# Pangenome mining of the *Streptomyces* genus redefines their biosynthetic potential

**DOI:** 10.1101/2024.02.20.581055

**Authors:** Omkar S. Mohite, Tue S. Jørgensen, Thomas Booth, Pep Charusanti, Patrick V. Phaneuf, Tilmann Weber, Bernhard O. Palsson

## Abstract

**Background:** *Streptomyces* is a highly diverse genus known for the production of secondary or specialized metabolites with a wide range of applications in the medical and agricultural industries. Several thousand complete or nearly-complete *Streptomyces* genome sequences are now available, affording the opportunity to deeply investigate the biosynthetic potential within these organisms and to advance natural product discovery initiatives.

**Result:** We performed pangenome analysis on 2,371 *Streptomyces* genomes, including approximately 1,200 complete assemblies. Employing a data-driven approach based on genome similarities, the *Streptomyces* genus was classified into 7 primary and 42 secondary MASH-clusters, forming the basis for a comprehensive pangenome mining. A refined workflow for grouping biosynthetic gene clusters (BGCs) redefined their diversity across different MASH-clusters. This workflow also reassigned 2,729 known BGC families to only 440 families, a reduction caused by inaccuracies in BGC boundary detections. When the genomic location of BGCs is included in the analysis, a conserved genomic structure (synteny) among BGCs becomes apparent within species and MASH-clusters. This synteny suggests that vertical inheritance is a major factor in the acquisition of new BGCs.

**Conclusion:** Our analysis of a genomic dataset at a scale of thousands of genomes refined predictions of BGC diversity using MASH-clusters as a basis for pangenome analysis. The observed conservation in the order of BGCs’ genomic locations showed that the BGCs are vertically inherited. The presented workflow and the in-depth analysis pave the way for large-scale pangenome investigations and enhance our understanding of the biosynthetic potential of the *Streptomyces* genus.

## Background

*Streptomyces*, a genus of soil bacteria, is known for its ability to produce various natural products that have applications in medicine and biotechnology. These organisms are characterized by their complex and diverse biosynthetic gene clusters (BGCs), which are responsible for the biosynthesis of these bioactive compounds [1]. Over the past decades, several genomic studies have revealed that the full range of metabolites produced by *Streptomyces* and the associated biosynthetic pathways are not yet fully known [2].

The same genomic studies have revealed extensive genomic and phylogenetic diversity within the Streptomyces genus. This diversity provides a huge potential for natural product discovery, but at the same time complicates comparative analyses across different species and strains. To mitigate this challenge, there is a growing consensus for the need to cluster *Streptomyces* into distinct groups or genus-equivalents [3,4]. Such refined classification aims to facilitate more precise comparisons to understand the biosynthetic diversity and evolution within the genus.

Recent advances in sequencing technology and genome mining tools have allowed for the data-driven discovery of natural products [5]. Several genome mining tools such as antiSMASH, BiG-SCAPE, and BiG-SLICE have revealed that various bacterial species encode previously unknown biosynthetic potential [6–9]. While genome mining tools have significantly advanced our understanding of biosynthetic potential, there is a recognition that the estimates of diversity and novelty can be constrained by the inherent limitations of these individual tools and reference databases. These limitations include inaccurate definitions of BGC boundaries or incomplete entries in reference databases such as MIBiG [10]. The strategy of integrating results from different tools can partially mitigate these challenges [11].

Large-scale pangenome mining studies help to understand the evolutionary patterns of biosynthetic gene clusters (BGCs) along with a deep characterization of the biosynthetic repertoire of a given bacterial species or genus [12–15]. Detailed comparative studies are now gathering evidence that vertical inheritance facilitates the diversification of BGCs more frequently than horizontal gene transfer [16]. Earlier pangenomic investigations of *Streptomyces*, examining 121 genomes [17] and 205 genomes [18], respectively, have underscored that the pangenome of *Streptomyces* is quite open and represents high diversity. These analyses brought to light a limited number of core genes—633 [17] and 304 [18]—found across all strains considered in each study, respectively. Recent sequencing efforts have significantly increased the publicly available high-quality genomes of *Streptomyces* [19]. In light of this explosion of sequencing data, there is an emerging need to re-investigate the *Streptomyces* pangenome and the biosynthetic diversity within these organisms.

In this study, we aim to address these questions by conducting the largest pangenome mining study of *Streptomyces* to date. By combining insights from various genome mining tools and clustering the organisms into distinct phylogroups, we seek to enhance our understanding of the biosynthetic potential, diversity, and evolutionary patterns inherent to this phylogenetically diverse genus.

## Results

### The dataset of *Streptomyces* genomes

In this study, we comprehensively analyzed genomes of the *Streptomyces* genus, sourcing both from the public database and from our newly published dataset [19]. As of 30 June 2023, we obtained accession IDs for 2,938 Streptomycetaceae genomes of all qualities from the NCBI RefSeq database (Data S1, Figure 1A). We also incorporated 902 newly sequenced [19] high-quality complete actinomycete genomes for a total of 3,840 genomes (Figure S1).

**Figure 1.**
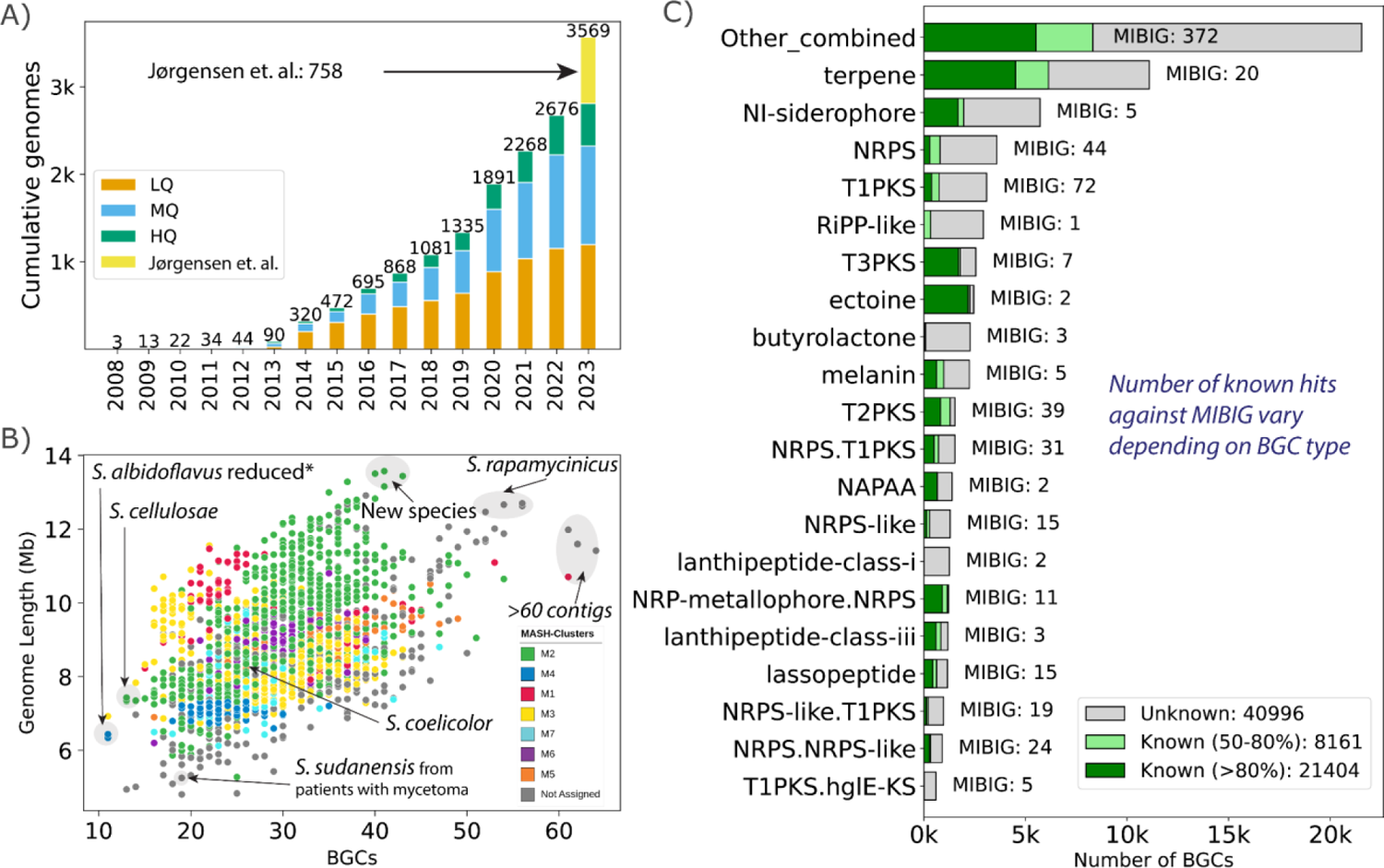
Dataset of *Streptomyces* genomes and BGC statistics. A) Number of *Streptomyces* genomes from the NCBI RefSeq database as of 30 June 2023. The final bar includes newly sequenced high quality genomes from our prior study [19]. Genomes are categorized by assembly quality: HQ (high-quality), MQ (medium-quality), and LQ (low-quality). B) Scatter plot illustrating the relationship between genome length and the number of BGCs in 2,371 genomes of the MQ and HQ categories. Annotations represent information on selected strains. C) Breakdown of the twenty most common types of BGCs detected in the HQ and MQ genomes. The remaining BGC types are bundled under “Other_combined,” which may contain hybrids of some of the listed types. Color-coded bars highlight BGC similarity percentages against the MIBiG database: gray for <50%, light green for 50-80%, and green for >80%. Bar annotations represent a tally of MIBiG entries with >80% similarity for the detected BGCs.

These 3,840 genomes were then curated further. To ensure a uniform taxonomic classification derived from whole genome sequences, we employed GTDB (version R214) [20,21] for taxonomic assignments (Figure S2). Out of the 3,840 genomes, 3,569 were identified as belonging to the *Streptomyces* genus. Using different assembly statistics, we grouped the selected 3,569 *Streptomyces* genomes into high-quality (HQ, 1,215 genomes), medium-quality (MQ, 1,156 genomes), and low-quality (LQ, 1,198 genomes) (Figure S1, S3). The final dataset, post-curation, included 2,371 genomes of sufficiently good quality (HQ or MQ).

We next classified the genomes at the species level. Of the 2,371 good-quality genomes, 1,956 were assigned to one of the 608 GTDB-defined species. The remaining 415 genomes lacked species assignments, representing potentially novel species beyond the GTDB catalog. Based on a 95% genomic similarity threshold using the MASH distance matrix, these 415 genomes were grouped into 202 species. Combining GTDB and MASH-based assignments, the dataset encompasses at least 810 *Streptomyces* predicted species, with 468 species represented by a single genome. Overall, these statistics indicate that the dataset is highly diverse, necessitating the careful grouping of these genomes for pangenome analysis.

The *Streptomyces* pangenome exhibited wide-ranging genomic characteristics. Genome sizes spanned from 4.8 Mbp to 13.6 Mbp, with a median of 8.5 Mbp (Figure 1B). Interestingly, the strains with the smallest genome sizes mainly belong to actinomycetoma-related pathogenic species of *S. sudanensis* and *S. somaliensis* [22]. In contrast, the largest-sized genomes primarily belong to *S. rapamycinicus* or to novel species. GC content ranged between 68.6% and 74.8%, with a median of 71.6%.

### Types of BGCs identified and similarity to known BGCs

Utilizing antiSMASH v7 [6], we identified a total of 70,561 BGCs in the 2,371 HQ or MQ genomes (Data S4). It is essential to highlight that genome quality can significantly influence the number of BGCs predicted for a particular genome. Specifically, when BGCs are located on contig edges, their count can be artificially increased when analyzed with antiSMASH as a broken BGC is likely to be counted twice. Thus the number of BGCs on the contig edge is a metric of genome quality for BGC analysis [23]. We identified only 6,524 BGCs (9.2%) situated at contig edges indicating a high quality of the collected dataset at capturing mostly complete BGCs [24]. Among the 1,215 genomes with complete assemblies (HQ), the number of BGCs per genome ranged between 11 and 56 with a median of 29 BGCs. It should be noted that the set of HQ assemblies included several *S. albidoflavus* strains in which multiple BGCs had been deleted, thus explaining the lower BGC count. The number of BGCs increased with the size of the genomes in accordance with prior observations (Figure 1B) [25,26].

The predominant BGC types in our dataset are terpene (11,095 BGCs), NRPS-independent (NI) siderophore (5,711 BGCs), nonribosomal peptide synthetase (NRPS) (3,599 BGCs), type1 polyketide synthase (T1PKS) (3,092 BGCs), ribosomally synthesized and post-translationally modified peptide like (RiPP-like) (2,933 BGCs), T3PKS (2,562 BGCs), ectoine (2,458 BGCs), butyrolactone (2,277 BGCs), melanin (2,244 BGCs), T2PKS (1,536 BGCs), and NRPS-T1PKS (1,536 BGCs). One can estimate the number of BGCs that encode known secondary metabolites by comparing the BGCs against the curated MIBiG database [10]. This estimate is provided automatically during antiSMASH analysis: the program generates “*knownclusterblast”* similarity scores that estimate how similar a certain region is to the BGCs in MIBiG by calculating a percentage of similar genes [6,10]. A threshold on the *knownclusterblast* score of greater than 80% of similar genes led to 21,404 BGCs (∼30%) that matched one of the 475 characterized BGCs from the MIBiG database. The most recurrent known BGCs were linked to the biosynthesis of compounds such as ectoine (2,230), desferrioxamine (1,685), geosmin (1,412), hopene (1,095), spore pigment (1,083), isorenieratene (852), albaflavenone (807), ε-Poly-L-lysine (730), and alkylresorcinol (708). These BGCs are known to be found commonly across the *Streptomyces* genus [27]. On average, 31% of the BGCs per genome matched to known BGCs in MIBiG. A further 8,161 BGCs (∼11.6%) had similarity scores between 60% to 80%, dominated by 1,116 hopene-like BGCs, while as many as 27,029 (38.3%) BGCs had similarity scores of less than 30%.

While estimates of novel BGCs provide valuable insights, they inherently depend on the completeness of the MIBiG database, potentially introducing bias. To further dissect this aspect, we examined the number of known BGCs across some of the abundant BGC types (Figure 1C). We found that certain BGC types—ectoine, NRP-metallophore-NRPS hybrid, T3PKS, T2PKS, lanthipeptide-class-iii, non-alpha polyamino group acids (NAPAA), and terpene—exhibited a significant similarity with the MIBiG database, as evidenced by over 40% of these BGCs having a *knownclusterblast* similarity of above 80%. In contrast, BGC types such as RiPP-like, lanthipeptide-class-i, butyrolactone, NRPS, NRPS-like, T1PKS, and NRPS-like-T1PKS hybrid showed less than 15% of their BGCs aligning with the MIBiG database with the same similarity threshold. However, it is essential to recognize that some BGC types, such as ectoine or NAPAA, are naturally less diverse and represent only a few compounds. For example, the majority of the ectoine-type BGCs (2,173 in total) were primarily aligned with just two MIBiG entries, both coding for the same compound ectoine (BGC0000853 and BGC0002052). Similarly, recognized BGC types like NAPAA, lanthipeptide-class-iii, melanin, and NI-siderophore matched fewer than eight MIBiG entries, and some of them are naturally less diverse.

### MASH-based analysis revealed 7 primary and 42 secondary MASH-clusters

Within the vast genomic landscape of the *Streptomyces* genus, the clearly defined classification of strains is essential for comparative analysis. Historically, comparative genome mining studies have predominantly centered on examining single species. However, this approach is limiting, especially in a genus like *Streptomyces* where many species are represented by a single genome sequence in public databases. This limitation has often necessitated a broader lens, encompassing genomes from the entire genus [4,17,18]. As valuable as genus-level insights are, the detection of over 800 species of *Streptomyces* demands a more focused approach. Accordingly, we sought to define distinct, sub-genus level groups.

Here, we propose a MASH-based whole genome similarity metric to empower comparative pangenome analysis by providing a statistical grouping of strains instead of the taxonomic delineations [28,29]. The MASH-clusters were generated by optimal K-means clustering in synergy with the highest average silhouette scores (Figure S4, Data S2). This analysis yielded seven primary MASH-clusters among 1,999 genomes, termed M1 through M7 (Figure 2, Data S2). To ensure the robustness of these clusters, a stringent silhouette score cutoff (0.4) was iteratively employed, leading to the removal of 372 genomes (Figure S5). These filtered genomes are less likely to be part of one of the 7 major MASH-clusters and may form additional clusters upon future sequencing efforts of these clades (Figure 2B). Venturing deeper, all primary MASH-clusters were subjected to an additional round of clustering, revealing 42 secondary MASH-clusters that encompassed 1,670 genomes after refinement based on silhouette scores with the same cutoffs (Figure S6-S12).

**Figure 2.**
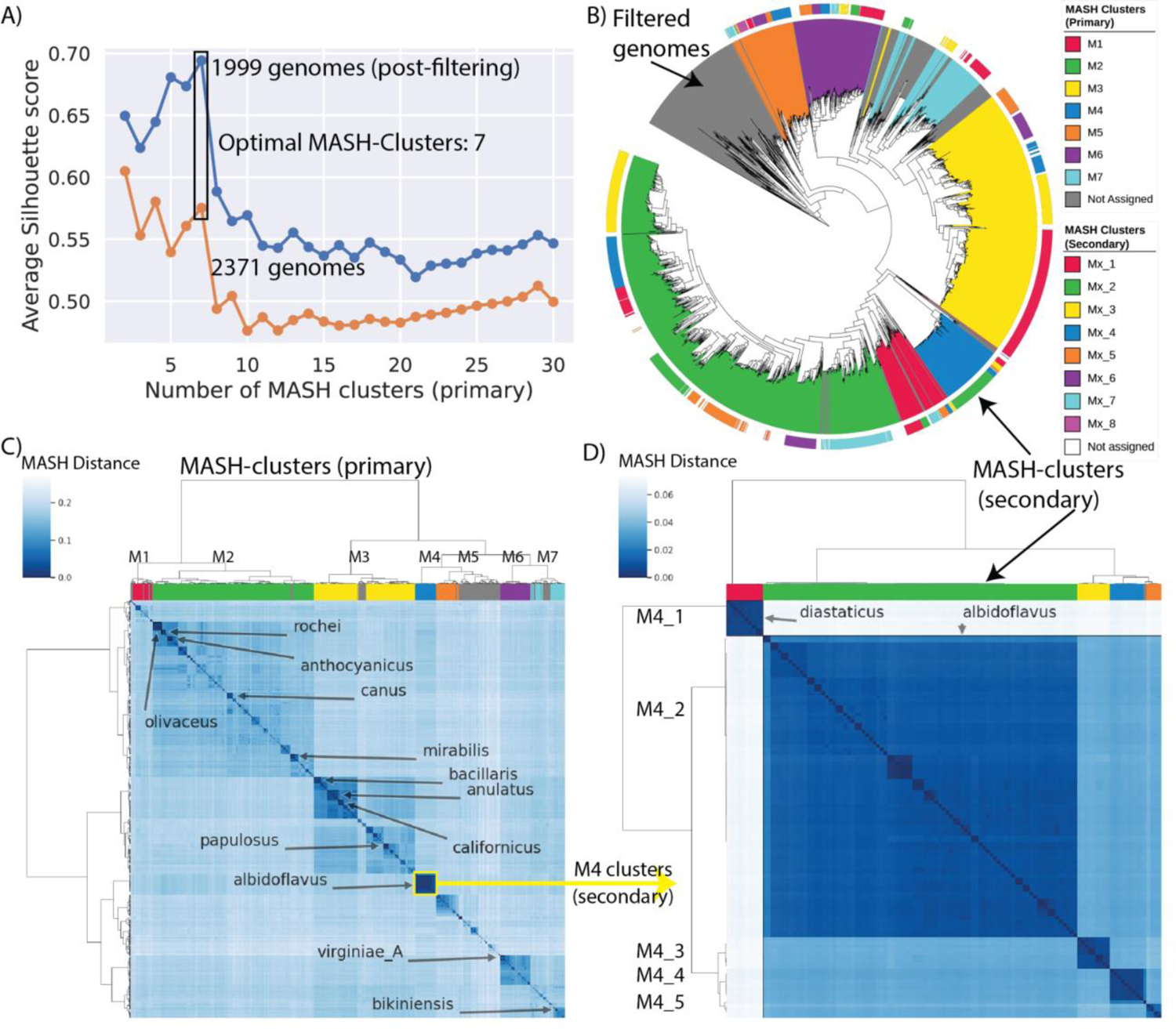
Mash-based clustering of the *Streptomyces* genus provides a basis for pangenome analysis. A) The average silhouette scores of all samples against the number of primary clusters with hierarchical clustering based on the MASH distance matrix. The orange line represents the original dataset of 2371 genomes, whereas the blue represents the dataset after filtering poorly clustered samples. B) A phylogenetic tree reconstructed using getphylo with *K. setae* strain KM-6054 as an outgroup. See Figure S13 for trees constructed using different methods and the consensus. The colored ranges represent the MASH-cluster assignment with gray color representing filtered genomes. The outer color strip represents the colors for secondary MASH-clusters (see Figures S6 to S12 for details). C) Heatmap representing the MASH distances between the 2371 genomes. The rows and columns are clustered using the hierarchical clustering method where the colors on columns represent the seven primary MASH-clusters (with gray color representing filtered-out genomes). The highlighted text on the heatmap represents some of the abundant species. D) Heatmap representing the MASH distances between the 119 genomes of the selected M4 cluster. The rows and columns are clustered using the hierarchical clustering method where the colors on columns represent the five secondary MASH-clusters (M4_1 to M4_5). M4_1 represents *S. diastaticus* whereas M4_2 to M4_5 represent different clusters within *S. albidoflavus*.

Several MASH-clusters stood out in this analysis. M2 emerged as the largest primary MASH-cluster representing 871 genomes. M2 harbors key species such as *S. coelicolor*, *S. rochei,* and *S. canus*. The second largest MASH-cluster, M3, represented 510 genomes with species such as *S. anulatus*, *S. bacillaris* and *S. papulosus*. MASH-cluster M5 stood out as the ancestral group and showed the poorest clustering (Figure 2C). However, a significant portion of genomes from MASH-cluster M5 were excluded from the refined MASH-clusters due to their low silhouette scores (Figure S5). MASH-cluster M4 represented 119 genomes, mostly of the species *S. albidoflavus (previously designated S. albus),* and was noteworthy for its high average clustering score (Figure 2D, Figure S9).

### Comparison of MASH-clusters with phylogenetic trees

The biggest drawback of using a similarity metric like MASH is the lack of an evolutionary model. Therefore, to evaluate the evolutionary relevance of the MASH-clusters, we compared them to genome-scale phylogenetic trees (Figure 2B). We constructed three trees by employing three distinct methodologies: autoMLST [30], GTDB-Tk (de novo workflow) [21], and getphylo [31] (Figure S13, Data S3). Upon comparison, a broad consensus was observed between the MASH-defined clusters and the clades delineated by different phylogenetic trees. There were, however, some outliers, chiefly clusters M1 and M7, which appeared to be paraphyletic (Figure S13). Upon closer inspection, however, these outliers fell within parts of the phylogenetic trees that were poorly supported and incongruent between the different methodologies. Further analysis revealed a striking level of incongruence between the three phylogenies. Only 62% of the branches were supported by a majority consensus and 33% by all three methodologies. The genus *Streptomyces* and its two major clades (represented by M2 and M3) are fully congruent, as well as many of the species and species complexes. However, lineages show a high degree of polytomy at the sub-generic level. This incongruence demonstrates the potential fallibility of phylogenetic methods when studying the intra-genus level relationships of *Streptomyces*.

Finally, we also compared the MASH-clusters with the RED_groups (relative evolutionary divergence-based groups) defined in a recent study as bacterial groups analogous to genera but characterized by equal evolutionary distance [4] (Figure S14). A consensus was observed for major groups except that the MASH-based method has split the RG_2 [4] into two separate MASH-clusters, M3 and M6. In this fashion, MASH-clustering proposed here complements the phylogenetic methods to produce statistically correlated groups.

### BGC diversity predictions based on known cluster similarity

To assess the diverse biosynthetic potential across genomes, it is helpful to group BGCs into gene cluster families (GCFs). GCFs are groups of BGCs that are homologous to each other, and thus are hypothesized to encode molecules that have similar chemical structures. GCFs are calculated by clustering BGCs using specialized tools such as BiG-SCAPE [7] or BiG-SLICE [8]. As a first step, we opted for BiG-SLICE (optimal with larger datasets) to execute this clustering across the entire dataset. Utilizing default parameters, we identified a total of 11,528 GCFs from the 70,561 BGCs (Figure 3A, Data S5). However, as highlighted in previous work [11], integrating diverse genome mining tools can enhance GCF refinement. For instance, minor genetic variations in regions adjacent to, but not directly involved in, biosynthesis can inadvertently lead to the classification of BGCs that code for identical secondary metabolites into disparate GCFs. To mitigate these issues, as a second step of defining GCFs, we used antiSMASH’s *knownclusterblast* results (with a similarity threshold of >80% of genes) to regroup the GCFs with the presence of predicted known BGCs.

**Figure 3.**
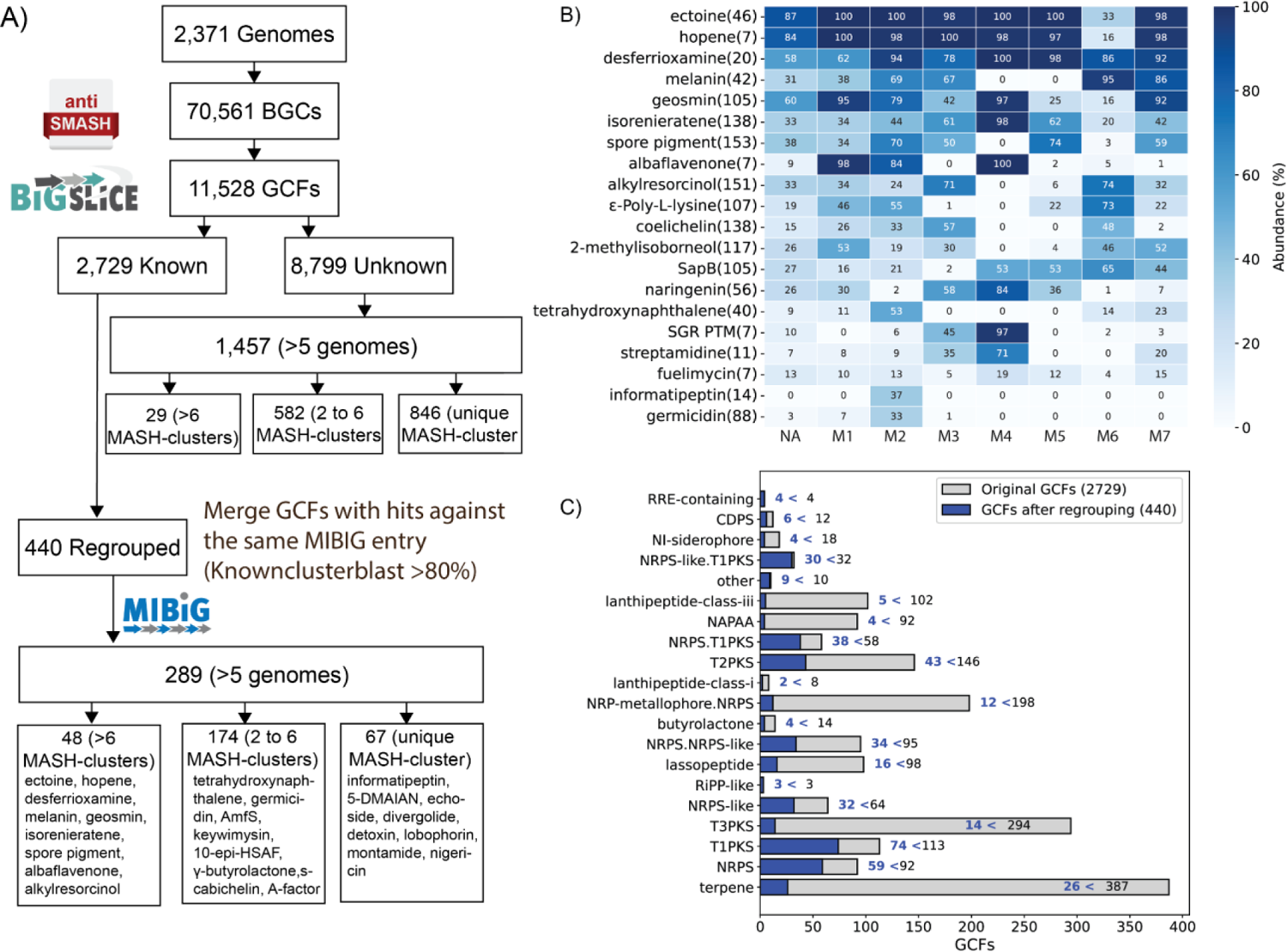
Advanced clustering of BGCs redefines known GCFs with reduced diversity in specific types of BGCs. A) Workflow used to detect BGCs, GCFs based on BiG-SLICE, and regrouping GCFs based on *knownclusterblast* similarity (>80% of genes). Several examples of known GCFs are reported in the bottom boxes, classified into common, accessory, or unique GCFs to MASH-clusters. B) Percentage abundance of the top twenty known GCFs across different primary MASH-clusters. Each row corresponds to a known compound (GCF). The number in parentheses denotes the number of BiG-SLICE detected GCFs that were regrouped into one GCF. C) Overview of the number of GCFs that were regrouped across the twenty most abundant BGC types. Gray bars represent the number of GCFs detected using only BiG-SLICE, whereas blue bars represent the reduced number of GCFs after regrouping based on *knownclusterblast*.

A total of 2,729 GCFs predicted by BiG-SLICE in the first step were associated with known secondary metabolites according to *knownclusterblast* results. After regrouping these GCFs at the second step, we effectively reduced the count of known GCFs from 2,729 to 440 (Figure 3A, Data S5). For instance, BGCs that code for spore pigment, alkylresorcinol, coelichelin, and isorenieratene (in the second step) were detected as 153, 151, 138, and 138 different GCFs (in the first step), respectively (Figure 3B). We investigated whether these reductions of GCF diversity predictions are dependent on the type of BGCs (Figure 3C). For example, BGC types such as lanthipeptide-class-iii, terpene, NRP_metallophore-NRPS hybrid, and T3PKS showed a high level of reduction in the diversity of GCFs when *knownclusterblast* results were integrated. In contrast, types such as NRPS or T1PKS showed a relatively lower reduction in the diversity of GCFs as was predicted in the first step using BiG-SLICE (Figure 3C).

We also note that these regrouped GCFs could contain minor internal variations. For a more precise investigation, we constructed a similarity network of two regrouped GCFs coding for spore pigment (153 originally predicted GCFs) and isorenieratene (138 originally predicted GCFs). We used a BiG-SCAPE generated distance matrix to create this network (more optimal for a relatively small dataset) (Figure S15A, and S15C). We also aligned the selected BGCs, which showed that the overestimated diversity of these known BGCs can be attributed to inaccurate BGC boundaries. For instance, variation in BGCs from different MASH-clusters was largely due to differences in the neighboring regions of the detected BGCs. The differential neighboring regions causing variation within the regrouped GCF were generally conserved within genomes from the same MASH-clusters (Figure S15B and S15D). In general, we observed that the types requiring fewer genes for core biosynthesis, such as terpene, T2PKS, T3PKS, siderophore, or RiPPs, were also among the most affected by these variations in neighboring regions.

### Diversity of GCFs across genomes from different MASH-clusters

Subsequently, we examined the distribution patterns of GCFs across the genomes delineated by the seven primary MASH-clusters to identify BGCs associated with specific MASH-clusters (Figure 3B). MASH-cluster M2 contained 2,606 GCFs that did not appear in any other MASH cluster. Similarly, MASH-clusters M3 and M6 contained 811 and 648 GCFs, respectively, that were specific to those MASH clusters (Figure S16). We also note that a total of 2,338 GCFs were specific to the 372 genomes that were dropped from the MASH-cluster definitions, and are likely to represent further diversity. It is imperative to note that MASH-clusters M2 and M3 constitute the most populous clades which may explain their apparent diversity of GCFs.

To gain deeper insights into the biosynthetic signatures of different MASH-clusters, we analyzed all GCFs containing at least five BGCs. This encompassed 289 known and 1,457 putatively novel GCFs. We found that 48 of the known GCFs (such as ectoine, hopene, desferrioxamine, etc.) displayed a widespread genomic distribution, being present in genomes across all MASH-clusters. We detected 174 known GCFs (such as germicidin, streptamidine, SGR-PTM, etc.) in the genomes across multiple, but not all, MASH-clusters. Finally, 67 of the known GCFs (such as informatipeptin, 5-DMAIAN, echoside, etc.) were specific to genomes from only one of the major MASH-clusters, representing the biosynthetic signatures of these groups of genomes (Figure S17). We also observed the same pattern of conservation of unknown GCFs in specific MASH-clusters (Figure S18). This observed presence of GCFs across MASH-clusters implies certain BGCs are likely to be found in certain MASH-clusters at primary or secondary levels (Figure S17-S18).

### Conservation of chromosomal synteny of BGCs

Finally, we present a novel workflow to capture BGC diversity by analyzing synteny within a MASH-cluster. The diversity of the BGCs and their functions is computationally predicted using similarity metrics and by visualization of the similarity networks (e.g., using BiG-SCAPE detected similarity scores). To explore the syntenic relationship between BGCs, we extended this network by adding edges between the BGCs that are neighbors on the chromosomes. Thus, all BGCs in any genome would be connected by edges in the order of presence on chromosomes.

As an example, we selected 49 complete genomes from MASH-cluster M4 that were further grouped into five secondary MASH-clusters (M4_1 to M4_5) (Figure 4A, Data S6). These strains primarily belonged to *S. albidoflavus* (M4_2 to M4_5) and *S. diastaticus* species. The resulting network of BGCs showed remarkable conservation of the order in which the BGCs have evolved on the chromosomal location (Figure 4B). We observed that different BGCs are either inserted or deleted from specific locations while maintaining the order of the seven commonly present BGCs across the M4 MASH-cluster genomes. We also observed that these differences are conserved within the secondary MASH-clusters (Figure 4B). This observation implicates the vertical inheritance of BGCs as strains evolve across different clades or groups.

**Figure 4.**
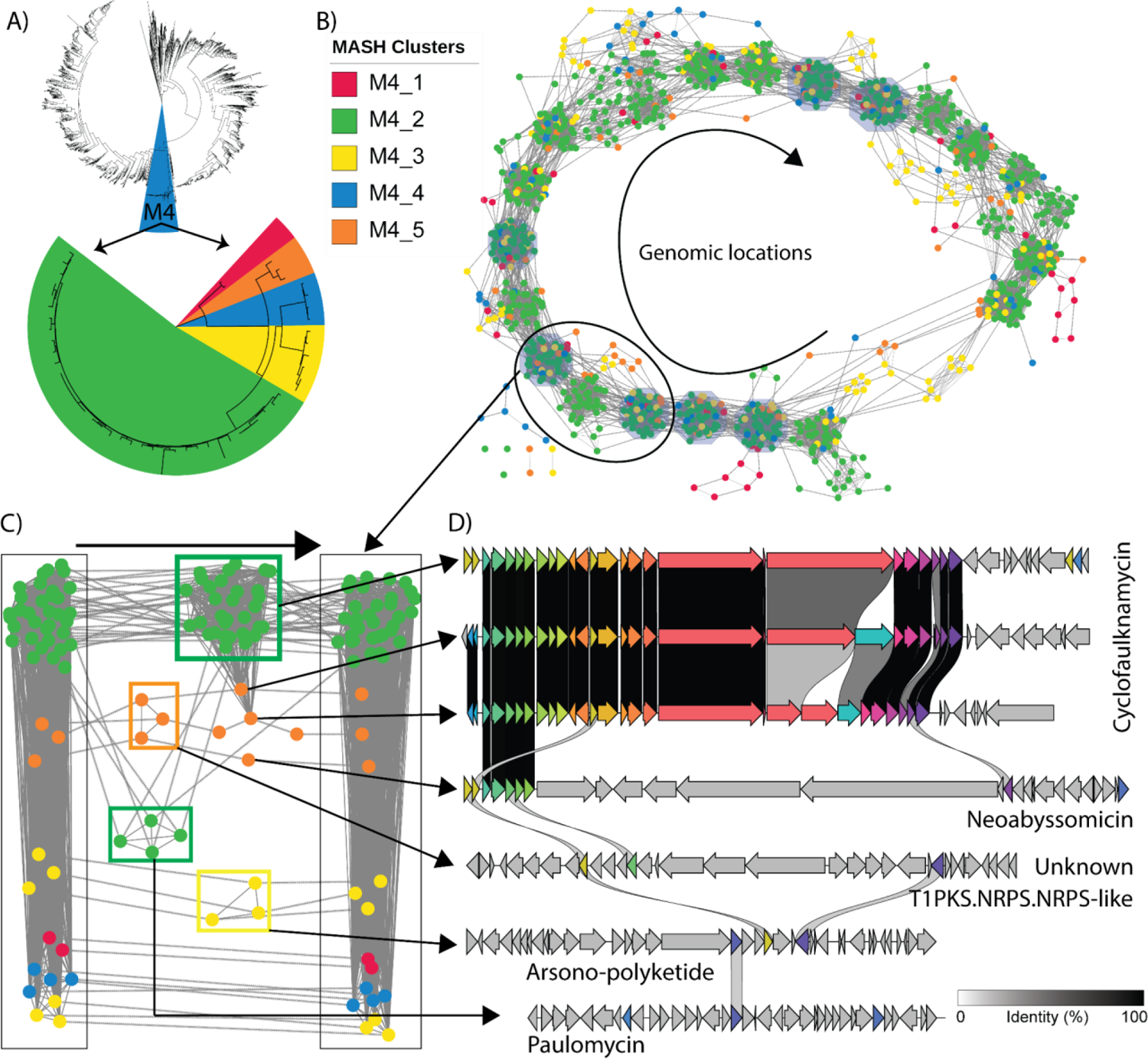
Synteny of BGCs across MASH-clusters M4_1 to M4_5 showed conserved and variable regions. A) Phylogenetic tree (top) of all 2371 genomes with highlighted M4 primary MASH-cluster. Phylogenetic tree (bottom) of complete HQ genomes from the M4 primary MASH-cluster grouped into five secondary MASH-clusters M4_1 through M4_5. M4_1 represents *S. diastaticus* whereas M4_2 to M4_5 represent different clusters within *S. albidoflavus*. B) Synteny network view of GCFs where the nodes represent detected BGCs across 49 high-quality complete genomes from M4. Seven of the BGCs were present across all 49 genomes and in the same order. The edges with solid lines represent BiG-SCAPE-based similarity between BGCs. C) A selected portion of the synteny network from part B. The leftmost BGC is a type 2 lanthipeptide and the rightmost BGC is a NI-siderophore. They are two of the seven BGCs conserved in all genomes. The middle BGCs are variable. D) Alignment of several variable BGCs from part C across strains from different secondary MASH-clusters.

We focused on a specific region between two of the conserved BGCs coding for a type 2 lanthipeptide and NI-siderophore (Figure 4C). The genomes belonging to MASH-clusters M4_1 and M4_4 did not possess any BGCs in this chromosomal region, along with some of the M4_3 genomes. The majority of the M4_2 genomes harbored an NRPS BGC coding for the known molecule cyclofaulknamycin (Figure 4D). The genomes of the M4_5 MASH-cluster showed interesting variation in this region. Two genomes were observed to harbor a reduced version of the cyclofaulknamycin BGC that could have a differential or loss of function, whereas the other M4_5 genome has acquired a completely different T1PKS BGC in the same region, one that codes for neoabyssomicin (Figure 4D)[32]. The genomes in M4_5 also harbor as yet uncharacterized T1PKS-NRPS hybrid BGC in the region. Some of the M4_2 genomes have additional BGCs in the region coding for the known PKS-like molecule paulomycin, whereas the others from M4_3 have a T1PKS-PKS-like hybrid that codes for arsono-polyketide, which is widespread across *Streptomyces sp.* [33].

To comprehend the actual diversity and investigate the evolution of BGCs, adopting a chromosomal syntenic perspective emerged as a crucial strategy. This recurrent pattern held true not only within MASH-cluster M4 but extended across our entire dataset, underscoring the robustness of the analytical framework. For instance, the incorporation of syntenic relationships into the similarity network revealed distinctive variations among five species within the M2_3 secondary MASH-cluster, including the model strain *S. coelicolor* A3(2) (Figure S19). This multi-dimensional overview of BGC diversity, grounded in expansive, high-quality genomic data, establishes a comprehensive tool for unraveling the nuanced intricacies of BGC evolution.

## Discussion

In this study, we conducted pangenome mining of the biosynthetic potential inherent in the *Streptomyces* genus, leveraging a dataset totaling over 2,370 genomes. Our investigative approach was underpinned by a comprehensive workflow that encompassed crucial steps for robust analysis. These steps included taxonomic identification, data quality checks, MASH-based clustering, as well as the detection of BGCs and GCFs. Furthermore, our methodology involved the regrouping of known GCFs to discern functional diversity and a thorough examination of synteny among BGCs distributed across the chromosomes. This comprehensive analytical framework has provided insights into both genomic architecture and the functional diversity inherent in these prolific secondary metabolite producers.

We emphasized the critical role of data curation as the foundational step in comparative genomic analysis, ensuring the establishment of a consistent dataset. Our workflow included an assessment of critical assembly metrics such as the number of contigs, N50 score, completeness, and contamination, enabling the classification of genomes into high, medium, and low quality. The genus’s vast diversity became evident, with as many as 810 detected species using GTDB and MASH. To delineate meaningful groups of genomes for comparative analysis, we employed a data-driven approach based on clustering MASH-based similarities. This methodology not only facilitated the grouping of genomes into distinct MASH-clusters but also emphasized those consistently clustered. While acknowledging the inherent limitations of clustering algorithms, we employed a strict silhouette score as a necessary metric, recognizing that these algorithms have their drawbacks, especially when dealing with unevenly distributed starting datasets. Consequently, we omitted 372 strains from MASH-cluster assignments, prioritizing the integrity of our analytical framework. Validation of the MASH-cluster definitions against different phylogenetic trees underscored the robustness of this grouping strategy for comparative analysis.

Expanding our analysis, the diversity and classification of GCFs were found to be notably influenced by several factors, including the type of BGC, the definition of BGC boundaries, and the completeness of the MIBiG database. This observation emphasizes the crucial role of manual inspection and refinement of existing genome mining tools in accurately characterizing the inherent diversity of detected BGCs. In the course of our study, the integration of similarity scores derived from *knownclusterblast* with the BiG-SLICE-based network highlighted a noteworthy finding—that the diversity of computationally predicted BGCs may be considerably constrained, especially in BGC types where the core biosynthetic regions are notably smaller than the predicted boundary regions. While we anticipate that improvements in GCF detection algorithms may yield more accurate predictions, the prediction of boundaries remains a substantial challenge in the genome mining field.

Leveraging the definition of MASH-clusters in our analysis, we identified GCFs demonstrating specificity or commonality across distinct MASH-clusters. Some of the common GCFs putatively coded for secondary metabolites such as ectoine, hopene, and desferrioxamine among others. This approach also facilitated the discernment of signature BGCs associated with groups of strains at different MASH-cluster levels. It is crucial to note that the variable size of MASH-clusters introduced variability in the number of signature BGCs observed across different clusters. As genome mining advances, these insights contribute to the ongoing refinement of methodologies, paving the way for more accurate and comprehensive assessments of biosynthetic potential across microbial genomes.

A detailed exploration of BGCs within MASH-cluster M4, which was further categorized into five secondary MASH-clusters, uncovered a striking observation - BGC order along the chromosome appears to be conserved. We observed shared genomic events such as deletions, insertions, and modifications of BGCs in specific chromosomal regions across distinct secondary MASH-clusters. Importantly, these patterns extend beyond M4, resonating across various MASH-clusters and species. Our investigation extends to the species level, exemplified by a comparative analysis involving five species, including *S. coelicolor*, within the secondary MASH-cluster M2_3 (Figure S19). This analysis illustrates how neighboring species have evolved distinct strategies to harbor diverse BGCs at specific chromosomal positions. The comparative examination provides insights into the evolutionary adaptations of these species, shaping their secondary metabolite biosynthetic capabilities. Notably, the findings underscore the role of vertical descent in the evolution of BGCs across species and MASH-clusters, aligning with a growing body of evidence in the literature [13,16,34,35].

With the exponential growth of genome sequencing, the influence of vertical descent is becoming increasingly apparent in the evolution of BGCs. The findings from this study significantly contribute to our understanding of these vertical inheritance mechanisms along with a need for manual inspection to more accurately capture the functional diversity of GCFs. These insights have broader implications for understanding the adaptive strategies employed by these prolific secondary metabolite producers in diverse ecological niches and environments.

## Conclusion

In conclusion, our study presents a pangenome analysis of the biosynthetic diversity of *Streptomyces*, a genus of high industrial importance. Data-driven clustering of nearly 2,400 *Streptomyces* genomes into MASH-clusters revealed 1) the diversity (or lack thereof) of computationally predicted BGCs, especially when automatically grouped into GCFs, 2) that certain BGCs/GCFs are specific to certain MASH-clusters, thus acting as potential biosynthetic signatures for the MASH-cluster, and 3) that synteny among BGCs are conserved, implying that vertical inheritance plays a major role in the evolution of BGCs. Taken together, our work not only contributes to advancing our understanding of secondary metabolite biosynthesis in *Streptomyces* but also highlights the evolving capabilities of pangenome analytics for biosynthetic diversity exploration.

## Methods

### Data collection, taxonomy detection, and quality check

The starting dataset to select *Streptomyces* genomes was gathered from two sources: NCBI and from those presented in a recent study [19]. As of 30 June 2023, we collected a total of 2,938 genomes of all assembly levels from NCBI RefSeq belonging to the family Streptomycetaceae (Data S1). We used this broader family of Streptomycetaceae with the aim of assigning taxonomy based on GTDB consistently (version R214) [20,21]. We collected an additional 902 of the 1,034 actinomycete genomes from a recent study [19] (Data S1). We note that 121 genomes of the 1,034 were already available on NCBI on 30 June 2023 and 11 were added later to the other study [19]. These genomes were processed through BGCFlow and different tools to assess the quality of the genomes were run [11]. The BGCFlow workflow used for the generation of results is available at https://github.com/NBChub/bgcflow. Out of these 3,840 genomes, 3,569 were identified as belonging to the *Streptomyces* genus as per GTDB definitions (Data S1, Figure S2).

The *Streptomyces* dataset of 3,569 genomes was processed with multiple quality checks. We calculated genome completeness and contamination metrics using CheckM [36]. When cutoffs of greater than 90% completeness and less than 5% contamination were used, 59 genomes were found to have low-quality assemblies (Figure S3). We also used the assembly statistics on the contigs and N50 scores for further curation. The genomes designated as complete or chromosome-level assembly as per NCBI were classified as high-quality (HQ). From the remaining genomes with scaffold or contig level assembly, we further annotated the genomes with more than 100 contigs or N50 score of less than 100 kb as low-quality (LQ). Genomes with fewer than 100 contigs were classified as medium-quality (MQ) (Figure S3).

In total, there were 1,215 HQ, 1,156 MQ, and 1,198 LQ genomes (Figure S1, S3). We defined “good quality” genomes as a set of 2,371 high-quality and medium-quality genomes. The date of submission was extracted from the NCBI metadata to represent the historical progress of genomes with different genome qualities (Figure 1A).

### MASH-based clustering analysis

The GTDB taxonomy assignment revealed 608 species for 1,956 genomes in the dataset of 2,371 curated *Streptomyces* genomes. The remaining 415 genomes lacked species assignments as they did not have similar representatives in the GTDB database. We calculated the MASH-based similarity network where the edges represent genome-wide similarity of greater than 95% (typical threshold for species detection). We used the community detection method [37] to define the best partitions that were assigned different MASH-based species totaling up to 202 novel species. Four species were highly represented (>30 genomes) in our dataset: *S. albidoflavus* (or *S. albus*) (109 genomes), *S. anulatus* (58 genomes), *S. olivaceus* (46 genomes) and *S. bacillaris* (33 genomes). Given such a diverse dataset, we proposed MASH-based clustering of the dataset as explained below.

We used a whole genome sequence similarity-based workflow to cluster the genomes into different subgroups of the *Streptomyces* genus. A similar workflow with MASH-based analysis was shown to capture the phylogroups in the past [29]. Following this method, we calculated MASH-distance for all pairs of genomes in the dataset using a BGCFlow rule that runs MASH (Data S2) [28]. We computed pairwise distances using Pearson’s correlation coefficient and performed hierarchical clustering using the ward.D2 method.

We added additional steps to the MASH-based analysis method [29] to identify the optimal number of clusters. We followed the elbow method to find the optimal number of k-means clusters and validated them using the average silhouette scores. We detected 7 optimal clusters based on both adjusted inertia for the K-means method and the high average silhouette score across the given dataset (Figure S4). The heatmap visualizations represented the diverse MASH-clusters defined here (Figure S4). Next, we visualized the silhouette scores across different MASH-clusters to validate the clustering using swarm plots (Figure S5). A random cutoff of 0.4 was chosen to select the genomes that have good cluster assignments. This cutoff results in the majority of the dataset being clustered consistently (except for MASH-cluster M5 that appears to be poorly clustered). We iteratively removed the poorly clustered genomes from the dataset until all genomes consistently scored above 0.4 on silhouette scores.

These curated steps resulted in the assignment of 1,999 genomes to a valid MASH-cluster (Data S2). We further identified MASH-clusters within each of the above-defined primary MASH-clusters. This secondary level of analysis led to the identification of 42 consistent secondary MASH-clusters across 1,670 of the 1,999 genomes. We note that the assignment of the MASH-clusters is dependent on the abundance of genomes collected in each cluster and will likely change as the number of genomes increases.

### Comparing MASH-clusters against phylogenetic trees

We constructed phylogenetic trees for all 2,371 curated Streptomyces genomes using an outgroup genome of the *Kitasatospora* genus (*K. setae* strain KM-6054). We used 3 different methods: GTDB-Tk [21], autoMLST [30], and getphylo [31] (Figure S13, Data S3). We calculated the consensus branch support depending on whether a particular branch was supported consistently by different methods (Figure S13). The branches supported by only one of the 3 trees were deleted to visualize the consensus tree (Figure S13D). We also extracted the RED_groups (relative evolutionary distance) calculated in a prior phylogenetic study based on GTDB [38]. These genomes were annotated on the color strip of the tree to compare against corresponding MASH-cluster assignments from our analysis (Figure S14). The tree visualizations were generated using iTOL [39]. The colored ranges for branches represented primary MASH-cluster assignments, whereas the outer colorstrip represent secondary level of MASH-cluster assignments. This qualitative comparison was used to compare MASH-cluster assignment results against the clades detected using different phylogenetic methods.

### Genome mining to detect BGCs

For a large-scale comparative pangenome analysis, it is common to annotate the genomes using a consistent method. Here, we annotated all genomes using prokka v1.14.6 [40]. We also used a list of seven selected genomes with high-quality manually curated annotations as a priority while running prokka using the parameter “*--proteins”* (Data S4). We used antiSMASH v7.0.0 on the annotated genomes to detect secondary metabolite BGCs (Data S4) [6]. The *knownclusterblast* results were used for primary assessment of whether the detected BGC regions show substantial similarity against the BGCs from MIBiG database [41]. We note that this parameter does have some pitfalls depending on various factors such as BGC region boundary definition, multiple BGCs being part of the same BGCs region, and incomplete information within the MIBiG database of some BGCs. Nonetheless, this metric still provides a way to quickly analyze large datasets such as the one presented here. We used a strict cutoff of greater than 80% *knownclusterblast* similarity to tentatively identify BGCs that produce known secondary metabolites (Figure 1C, Data S4). BGCs with 50 to 80% similarity were similarly marked as producers of known secondary metabolites but with lower confidence.

### Detection of GCFs

We detected over 70,000 BGCs. Subsequently, we used BiG-SLICE to calculate gene cluster families (GCFs) using the default parameters (threshold of 900) [8] (Data S5). We further annotated the GCFs as known if they had BGCs with *knownclusterblast* similarity above 80% (Figure 3A). Different GCFs that contained BGCs with hits against the same MIBiG entry were combined into a single “regrouped” GCF and putatively associated with known BGCs (Data S5). The regrouped GCFs may still represent minor functional diversity; however, the large number of GCFs often stemmed from variation in neighboring genes that were not part of the BGC as reported in MIBiG. For example, spore pigment, alkylresorcinol, coelichelin, and isorenieratene BGCs were regrouped from a large number of predicted GCFs. This study prioritized the analysis of the known BGCs and left the unknown BGCs out of such regrouping analysis.

The abundance of some of the common GCFs (after regrouping) was calculated across different MASH-clusters (Figure 3B). The UpSet plot was used to visualize the overlap of GCFs across MASH-clusters (Figure S16). Selected BGCs from two GCFs putatively coding for spore pigment and isorenieratene were further extracted for in-depth comparison. For more accurate similarity calculation, we used BiG-SCAPE to generate a similarity network with a default threshold of 0.3 on the distance metric [7]. The network was visualized using Cytoscape where node colors represented different MASH-clusters [42]. Representative BGCs from different BiG-SCAPE predicted GCFs were further chosen to visualize the BGC region alignment using clinker tool [43] (Figure S15).

### Integrated network of BGCs similarity and chromosomal order

We developed a custom workflow to simultaneously visualize BGC diversity and the order of BGCs along the chromosome. As a case study, we selected BGCs from 49 high-quality complete genomes from MASH-cluster M4 that spanned 5 secondary-level MASH-clusters (Figure 4, Data S6). Each node in the network represents a BGC, and nodes were connected with two types of edges. The first type represented BiG-SCAPE-based similarity. The second type reflected the order of BGCs present on the chromosome (Figure 4). A specific region of the chromosome with two conserved BGCs was extracted for manual inspection of the variation of this region (Figure 4C). The selected BGCs were visualized using the clinker [43] to observe the alignments (Figure 4D). A similar integrated network was also reconstructed for 23 genomes from 5 different GTDB-defined species that belonged to MASH-cluster M2_3 (Figure S19, Data S8).

## Declarations

### Ethics approval and consent to participate

Not applicable

### Consent for publication

Not applicable

### Availability of data and materials

All the data is available as supplementary materials in files Data S1 to Data S6. Data S1 contains the accessions of all the genomes used in the study.

## Authors’ contributions

B.O.P and O.S.M conceived the study; O.M., T.S.J., T.W., and B.O.P. designed the research; O.M. performed data analysis, implemented the workflow, and gathered the results; O.M., T.S.J, T.B. analyzed and interpreted the data, with assistance from P.C, P.V.P, T.W, and B.O.P; O.M., T.S.J, T.B., P.C., P.V.P., T.W., and, B.O.P. wrote and revised the article.

## Acknowledgments

We acknowledge Matin Nuhamunada for his guidance on the use of BGCFlow to enable data processing. We thank Emre Özdemir, Marc Abrams, Daniel Zielinski, Kai Blin, and Simon Shaw for valuable discussions.

## Supplementary Information for

### Legends for Data S1 to S6

**Data S1.** Metadata of the genomes used

**Genomes Metadata:** List of 3,840 genomes used at the start of the study with information on the source, quality assigned, taxonomic information based on GTDB, checkM metrics of assembly quality, bioproject accession numbers for all genomes, assigned MASH clusters at two levels for selcted *Streptomyces* genomes, and the RED groups from prior study.

**Data S2.** MASH-based clustering and silhouette scores

MASH distances: Table with MASH based distances across all 2,371 selected genomes.

Silhouette scores (primary): Table with list of 2,371 genomes with assigned clusters based on clustering analysis. The columns represent the assigned clusters after each filtering round (upto 5). The genomes being removed based on silhouette score cutoff of 0.4 are annotated as “Dropped”. The columns also mention the silhouette score at each round of filtering. The final round includes 1,999 genomes with primary MASH-cluster assignments.

Silhouette scores (secondary): Table with list of 1,999 genomes with assigned clusters at secondary level based on clustering analysis. The columns represent the assigned clusters after each filtering round (upto 3). The genomes being removed based on silhouette score cutoff of 0.4 are annotated as “Dropped”. The columns also mention the silhouette score at each round of filtering. The final round includes 1,670 genomes with secondary MASH-cluster assignments.

**Data S3.** Phylogenetic trees using different methods

Three different phylogenetic tree files were calculated using autoMLST, getphylo and GTDB-Tk methods. The iTOL project with all the phylogenetic tree can be found at the link below: https://itol.embl.de/shared/omkar31

**Data S4.** Detected BGCs across 2,371 genomes of HQ and MQ quality

BGCs counts: Table with number of BGCs detected in each of the genomes analyzed.

**BGC information:** Metadata table with information on each of the detected BGCs

**Data S5:** Detected GCFs using BiGSLiCE and regrouping of known GCFs

GCFs (BiGSLICE): List of detected GCFs using BiGSLICE with metadata on number of BGCs and combined GCF ID that were regrouped based on shared known clusters blast hits

GCFs (Regrouped): List of GCFs as defined in this study using BiGSLICE along with knownclusterblast similarity with metadata on number of BGCs BiGSLICE defined GCFs.

BGC information: Assignment of GCFs and combined GCFs for each BGC

**Data S6:** Cytoscape file of BiG-SCAPE similarity network for BGCs in M4 Mash-cluster

The network visualizations corresponding to Figures 4B and 4C. The network included edges based on BiGSCAPE similarity. Additional edges were added if the BGCs appeared next to each other on the chromosome.

---

**Figure S1.**
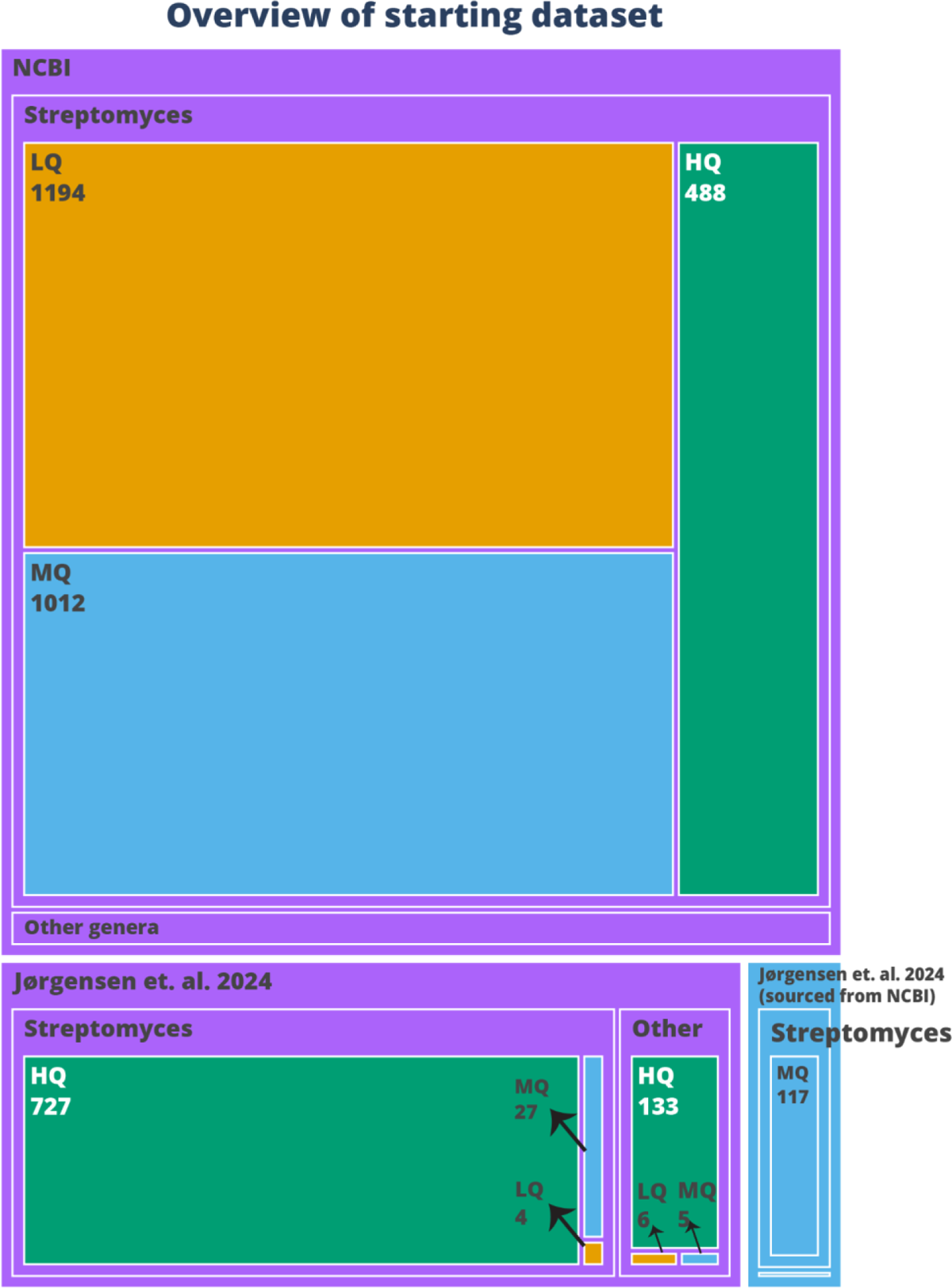
Dataset overview Treemap illustrating the number of genomes during various filtering stages. The primary rectangles denote the genome source. Genomes were sourced from NCBI on 30 June, 2023 and our prior study [1]. Part of the genomes form our prior study were already available at NCBI on 30 June, 2023 and were sourced from there. The secondary layer signifies the GTDB-based genus assignment to *Streptomyces*. The tertiary layer classifies genomes by the assembly quality as defined in this study: HQ (High Quality), MQ (Medium Quality), or LQ (Low Quality). HQ: Genomes with complete or chromosome-level assemblies. MQ: Genomes with contig or scaffold level assembly with less than 100 contigs. LQ: Genomes with contig or scaffold level assembly with more than 100 contigs.

**Figure S2.**
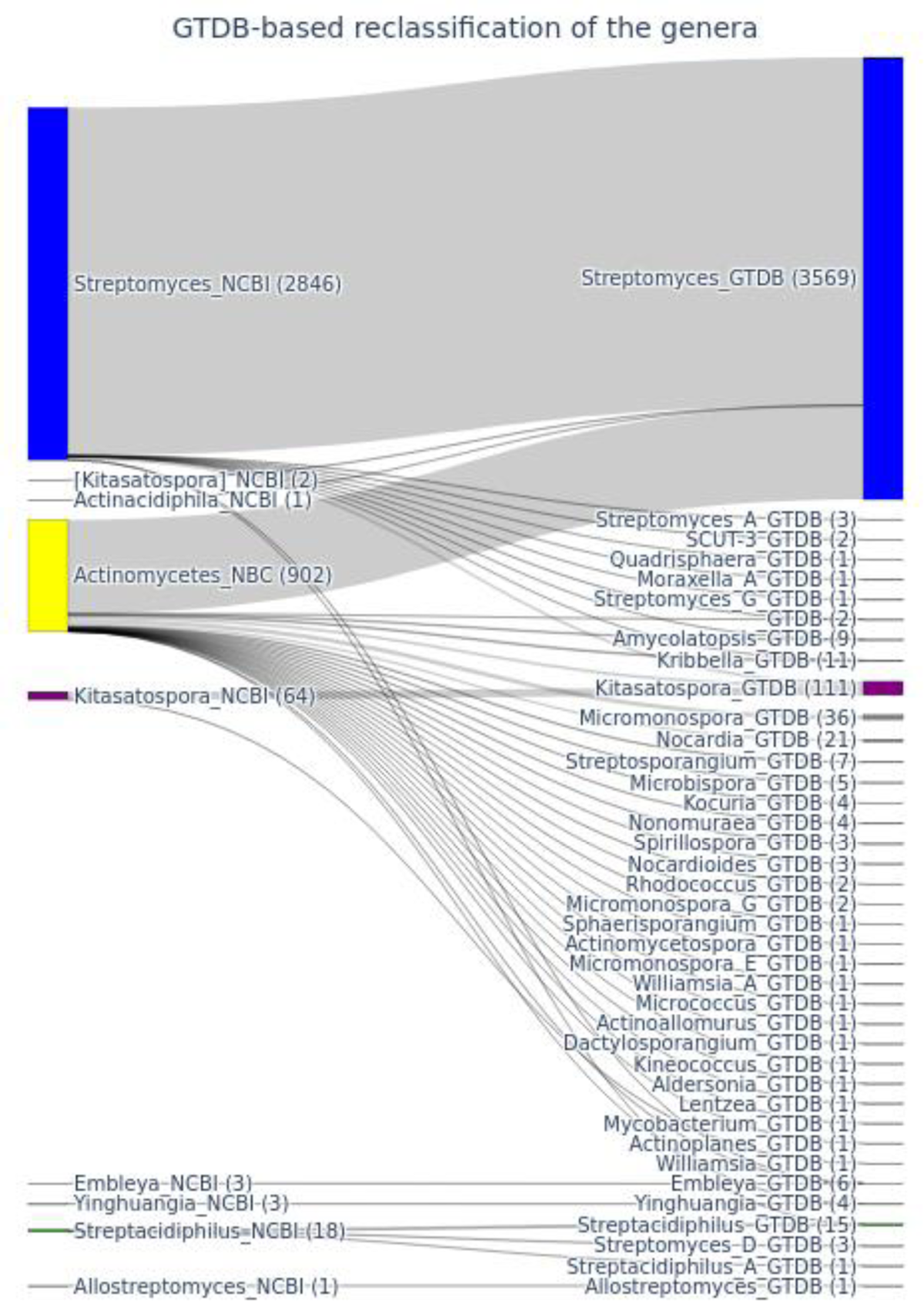
GTDB-based taxonomic assignment The genomes of the Streptomycetaceae family from NCBI RefSeq and actinomycetes from our recnet study (Jorgensen et al. 2024) (also known as NBC collection) (left) were assigned genus definitions based on GTDB R214 (right). Note that 38 *Streptomyces* genomes were reassigned to different genera using GTDB taxonomy (25 to *Kitasatospora*)

**Figure S3.**
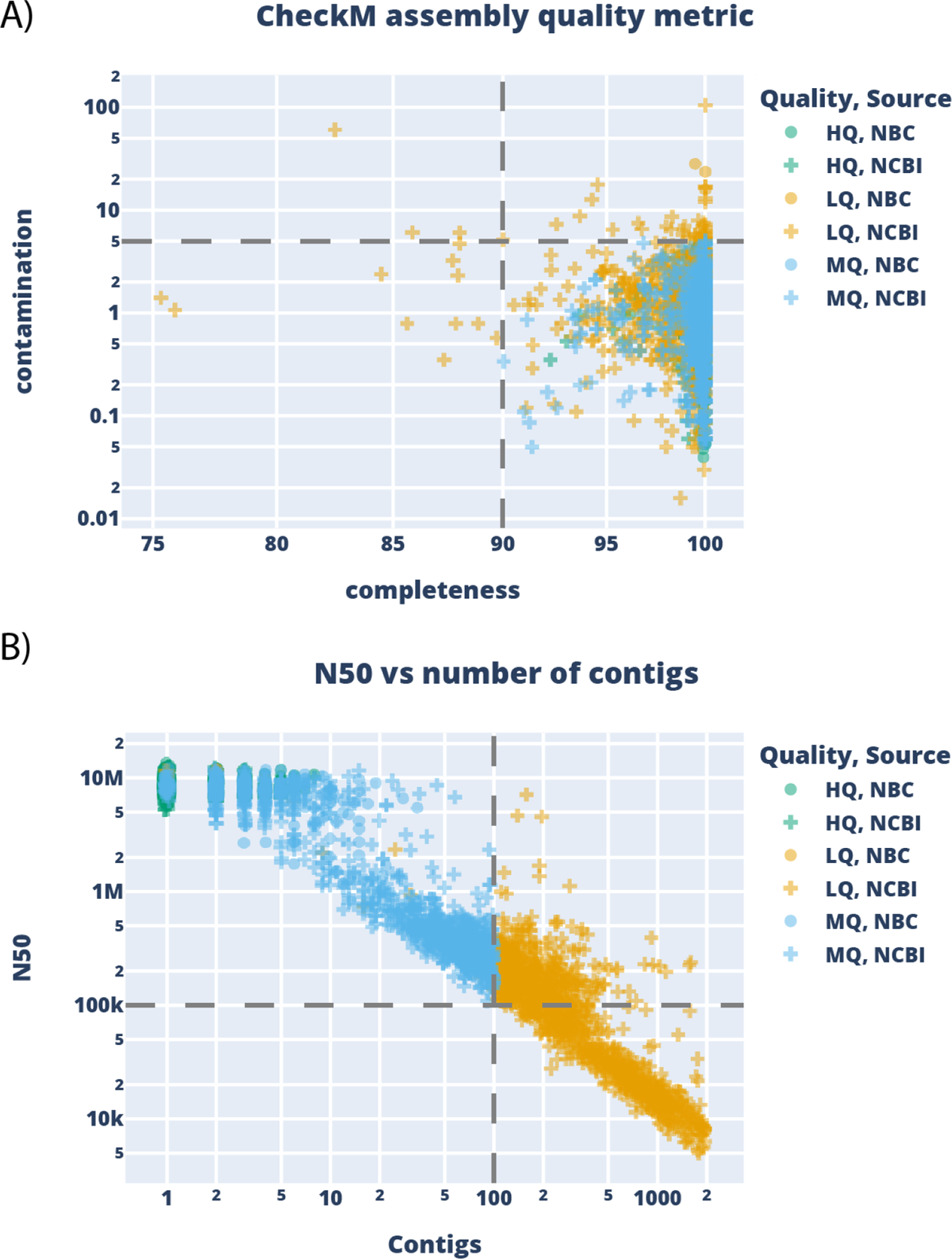
Assembly quality overview and filtering of the dataset A) Scatter plot representing the distribution of completeness and contamination score calculated using CheckM across 3,569 *Streptomyces* genomes. Genomes with completeness of less than 90% or contamination of more than 5% were dropped. B) Scatter plot representing the distribution of N50 score and number of contigs across 3,569 *Streptomyces* genomes. The colors represent the quality (HQ, MQ, or LQ) whereas the shapes represent the source of the genome (NCBI or NBC). HQ: Genomes with complete or chromosome level assemblies. MQ: Genomes with contig or scaffold level assembly with less than 100 contigs. LQ: Genomes with contig or scaffold level assembly with more than 100 contigs.

**Figure S4.**
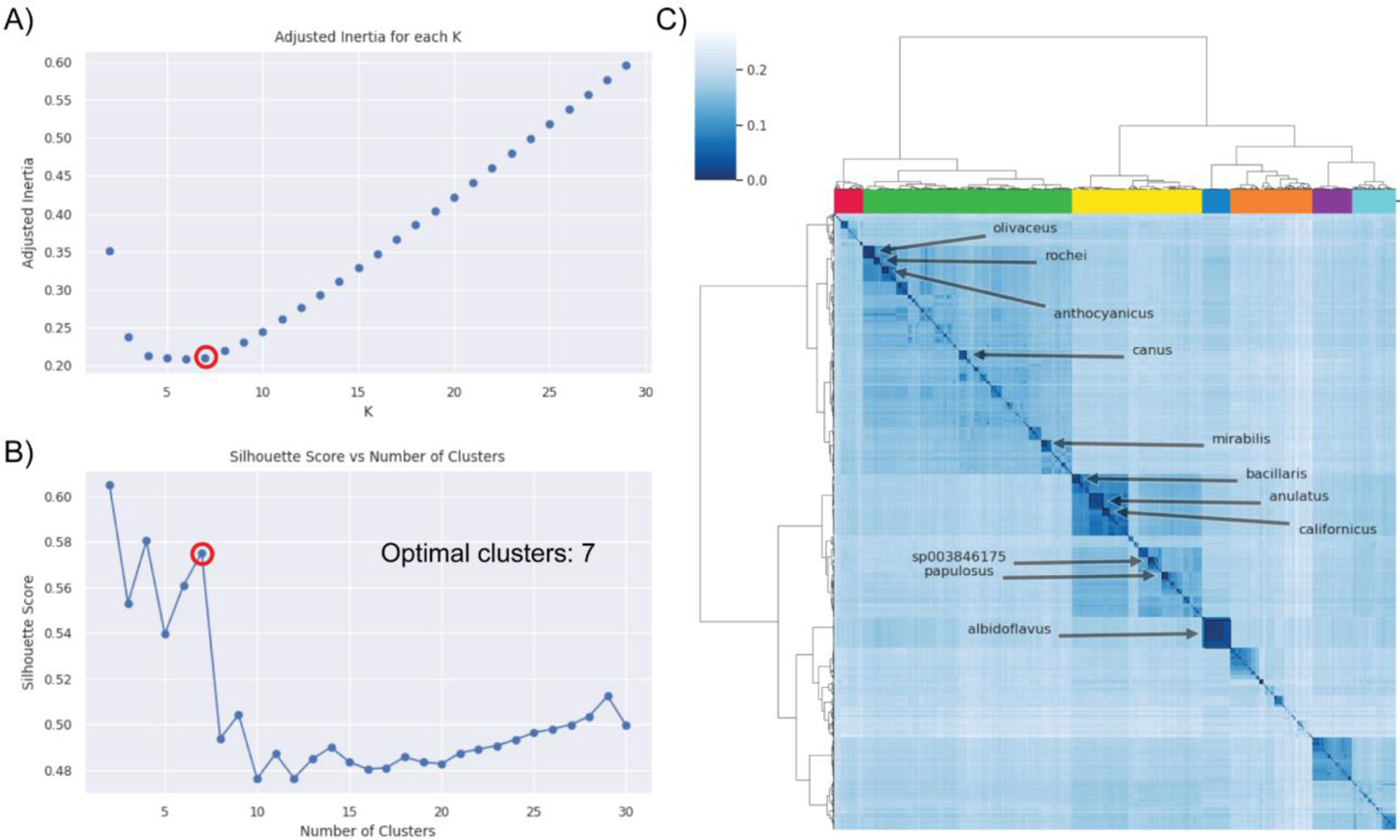
Detection of optimal clusters using K-means and Silhouette scores on the curated dataset of 2371 genomes A) Adjusted inertia against different K-means clusters representing optimal clustering with 7 Mash-clusters. B) Average Silhouette score of all samples for different numbers of clusters showing 7 optimal Mash-clusters. C) Hierarchical dendrogram with clustermap representing MASH distance values across genomes. The column colors represent the 7 optimal Mash-clusters. The top 20 abundant species are highlighted in the clustermap.

**Figure S5.**
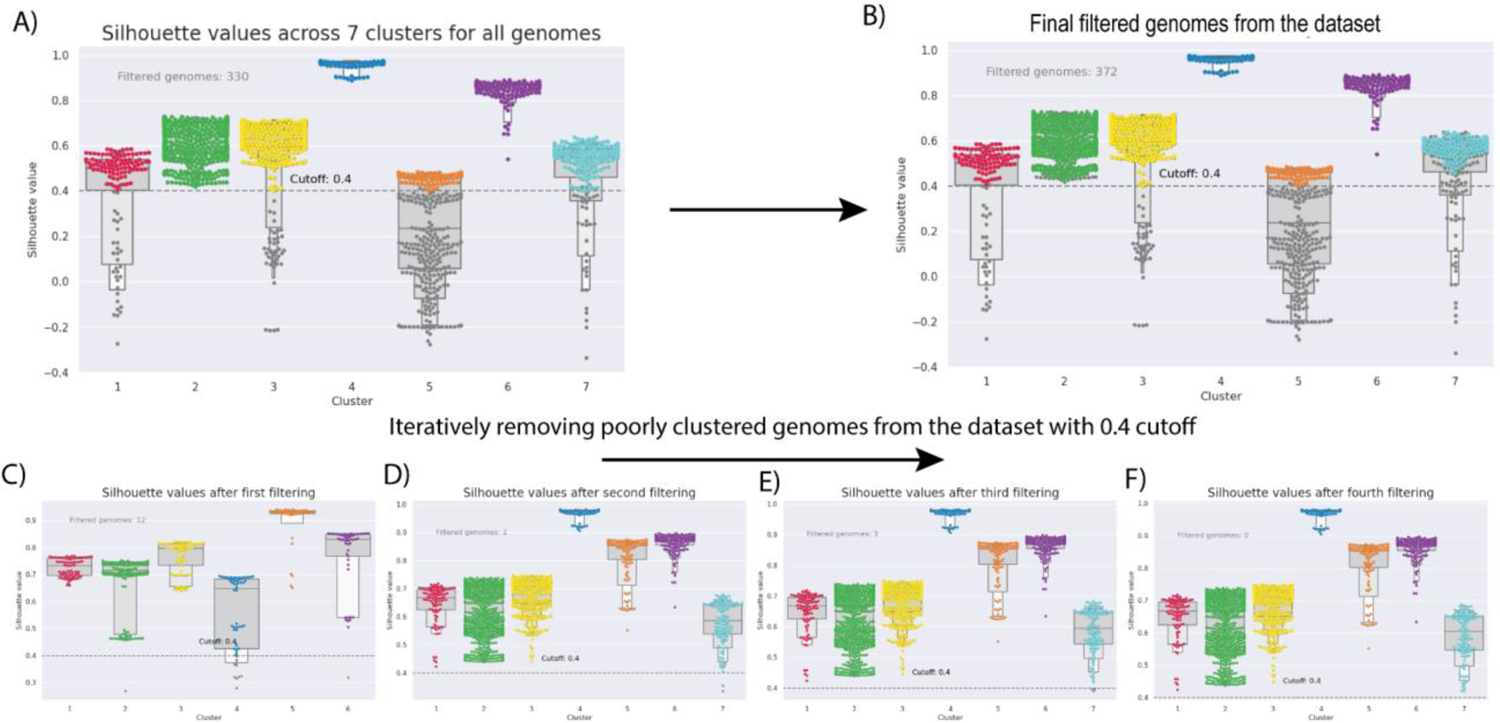
Iterative filtering of poorly clustered genomes using Silhouette score cutoff A) Swarmplot representing silhouette values of each sample genome across the 7 predicted MASH-clusters. The genomes with silhouette values lower than 0.4 are removed to detect the clusters accurately. C) to F) Iteratively reducing the size of the dataset until all samples have silhouette values higher than 0.4. B) Final dataset of 1999 genome samples plotted on the original clustering in panel A. The color of the dots represents one of the seven predicted primary MASH-clusters with grey color denoting the poorly clustered sample genomes that are filtered out.

**Figure S6.**
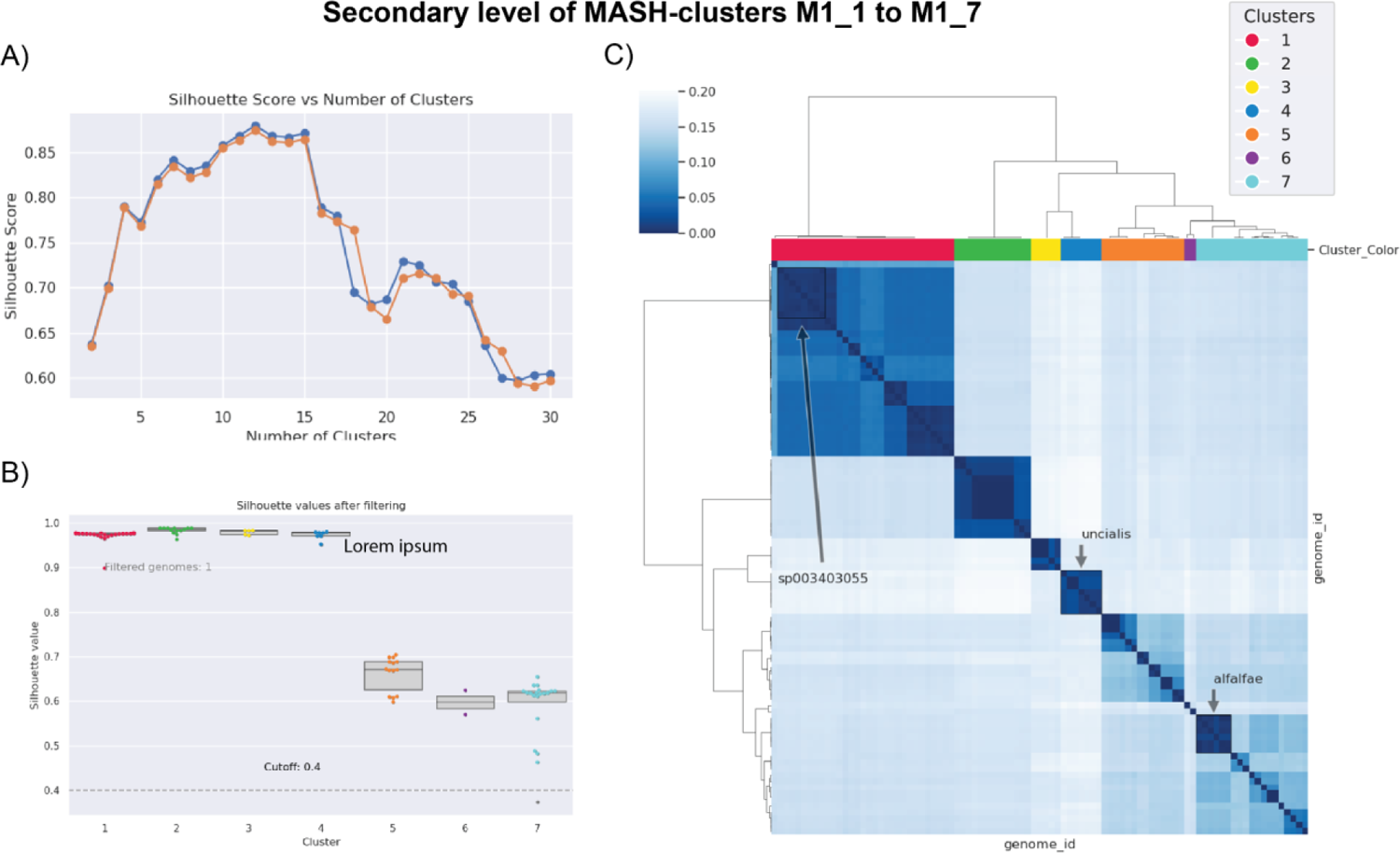
Detection of secondary MASH-clusters using Silhouette scores within the M1 primary MASH-cluster A) The average silhouette scores of all samples against the number of defined clusters with hierarchical clustering based on the MASH distance matrix. The orange line plot represents the original dataset of M1 MASH-cluster genomes whereas the blue represents the dataset after removing poorly clustered samples. B) The silhouette scores of each sample across 7 secondary MASH-clusters. The cutoff of 0.4 was used to select the samples with good clustering. The grey dots represent 1 genome that was removed from the clustering analysis. C) Heatmap representing the MASH distances between the genomes from the refined dataset. The rows and columns are clustered using the hierarchical clustering method where the colors on columns represent the 7 secondary MASH-clusters. The highlighted text on the heatmap represents some of the abundant species.

**Figure S7.**
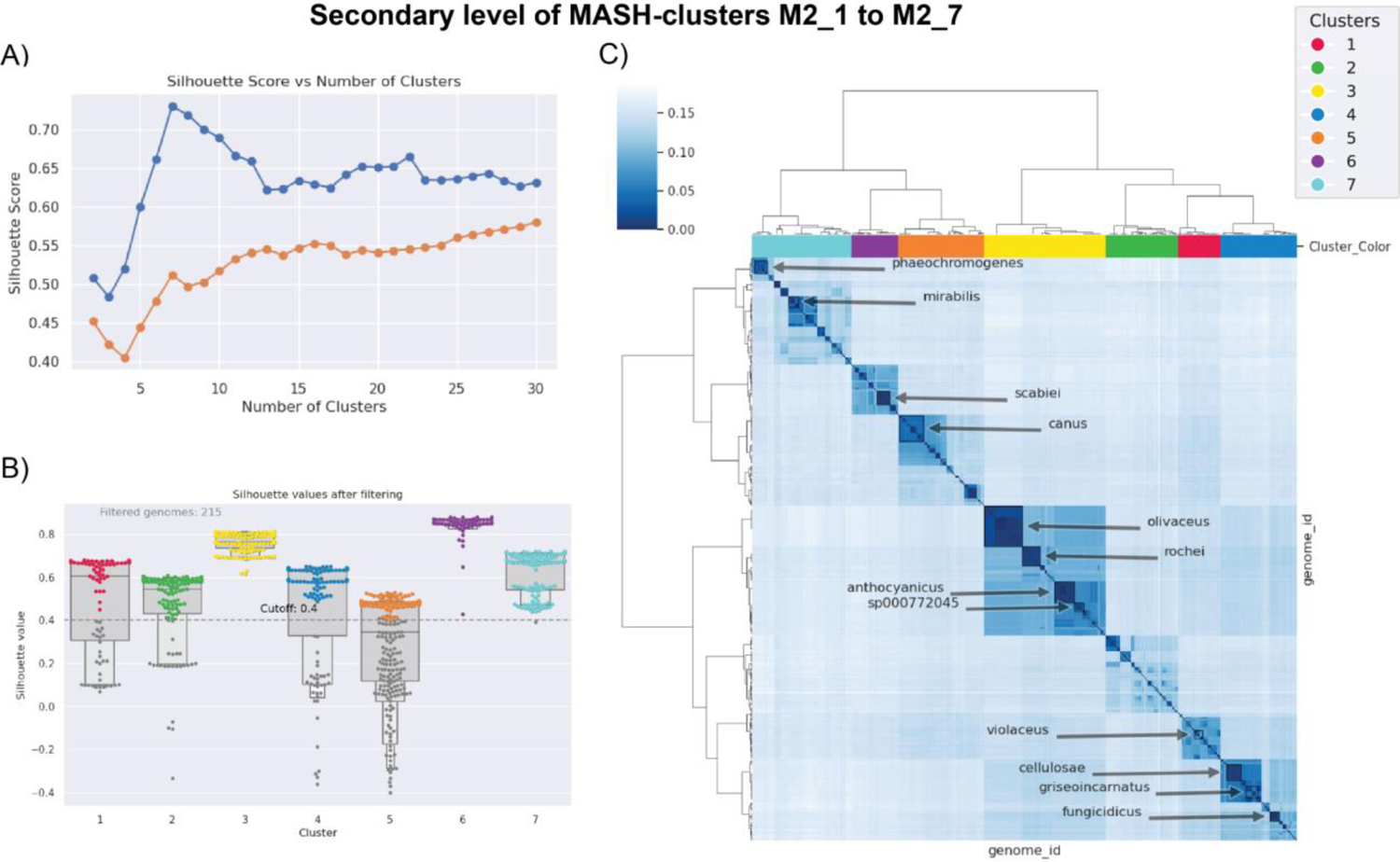
Detection of secondary MASH-clusters using Silhouette scores within the M2 primary MASH-cluster A) The average silhouette scores of all samples against the number of defined clusters with hierarchical clustering based on the MASH distance matrix. The orange line plot represents the original dataset of M2MASH-cluster genomes whereas the blue represents the dataset after removing poorly clustered samples. B) The silhouette scores of each sample across 7 secondary MASH-clusters. The cutoff of 0.4 was used to select the samples with good clustering. The grey dots represent 215 genomes that were removed from the clustering analysis. C) Heatmap representing the MASH distances between the genomes from the refined dataset. The rows and columns are clustered using the hierarchical clustering method where the colors on columns represent the 7 secondary MASH-clusters. The highlighted text on the heatmap represents some of the abundant species. Note: *S. anthocyanicus* is the renamed species of *S. coelicolor* as per GTDB.

**Figure S8.**
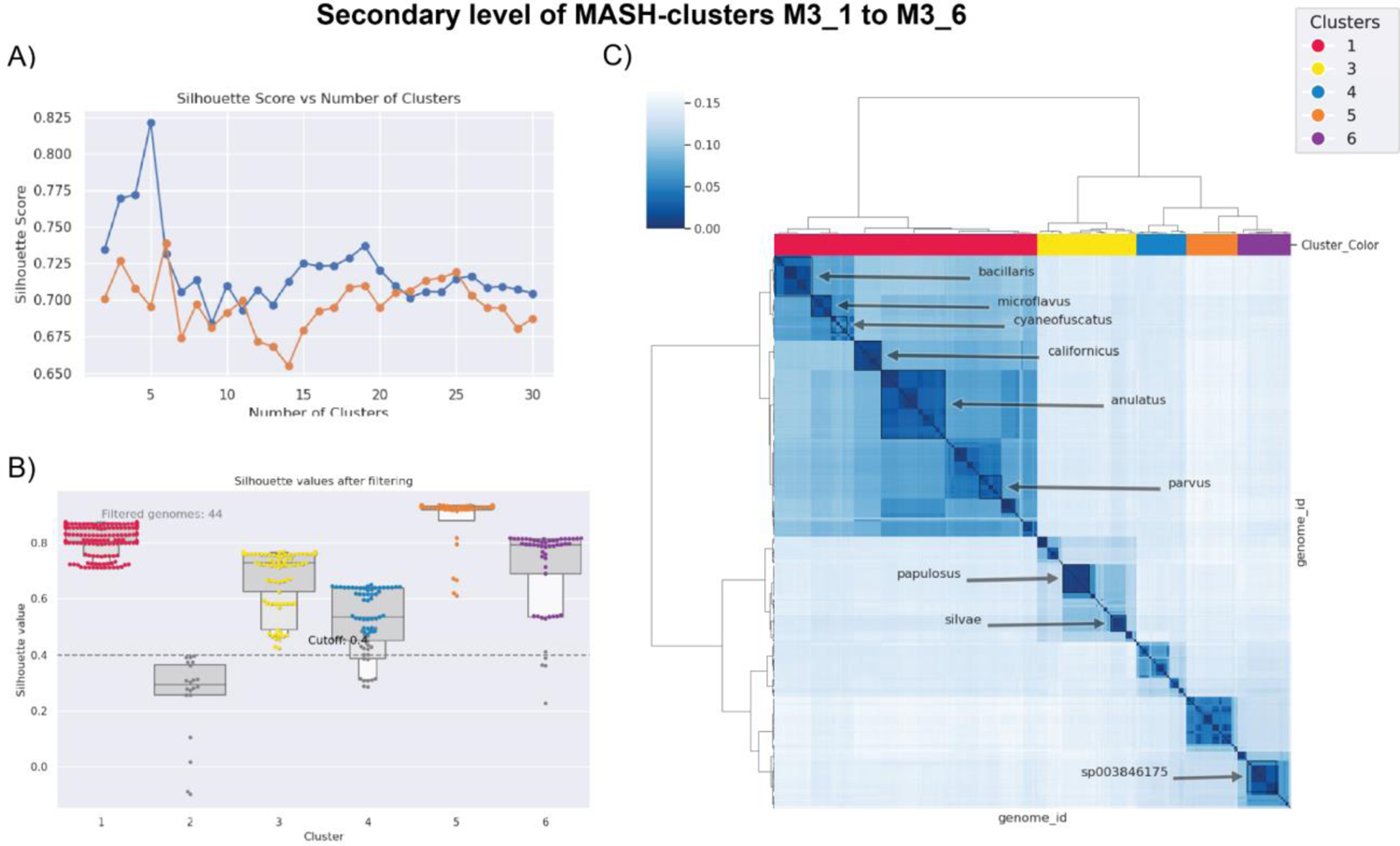
Detection of secondary MASH-clusters using Silhouette scores within the M3 primary MASH-cluster A) The average silhouette scores of all samples against the number of defined clusters with hierarchical clustering based on the MASH distance matrix. The orange line plot represents the original dataset of M3 MASH-cluster genomes whereas the blue represents the dataset after removing poorly clustered samples. B) The silhouette scores of each sample across 6 secondary MASH-clusters. The cutoff of 0.4 was used to select the samples with good clustering. The grey dots represent 44 genomes that were removed from the clustering analysis, including an entire cluster 2. C) Heatmap representing the MASH distances between the genomes from the refined dataset. The rows and columns are clustered using the hierarchical clustering method where the colors on columns represent the 5 secondary MASH-clusters (note that cluster 2 was completely removed). The highlighted text on the heatmap represents some of the abundant species.

**Figure S9.**
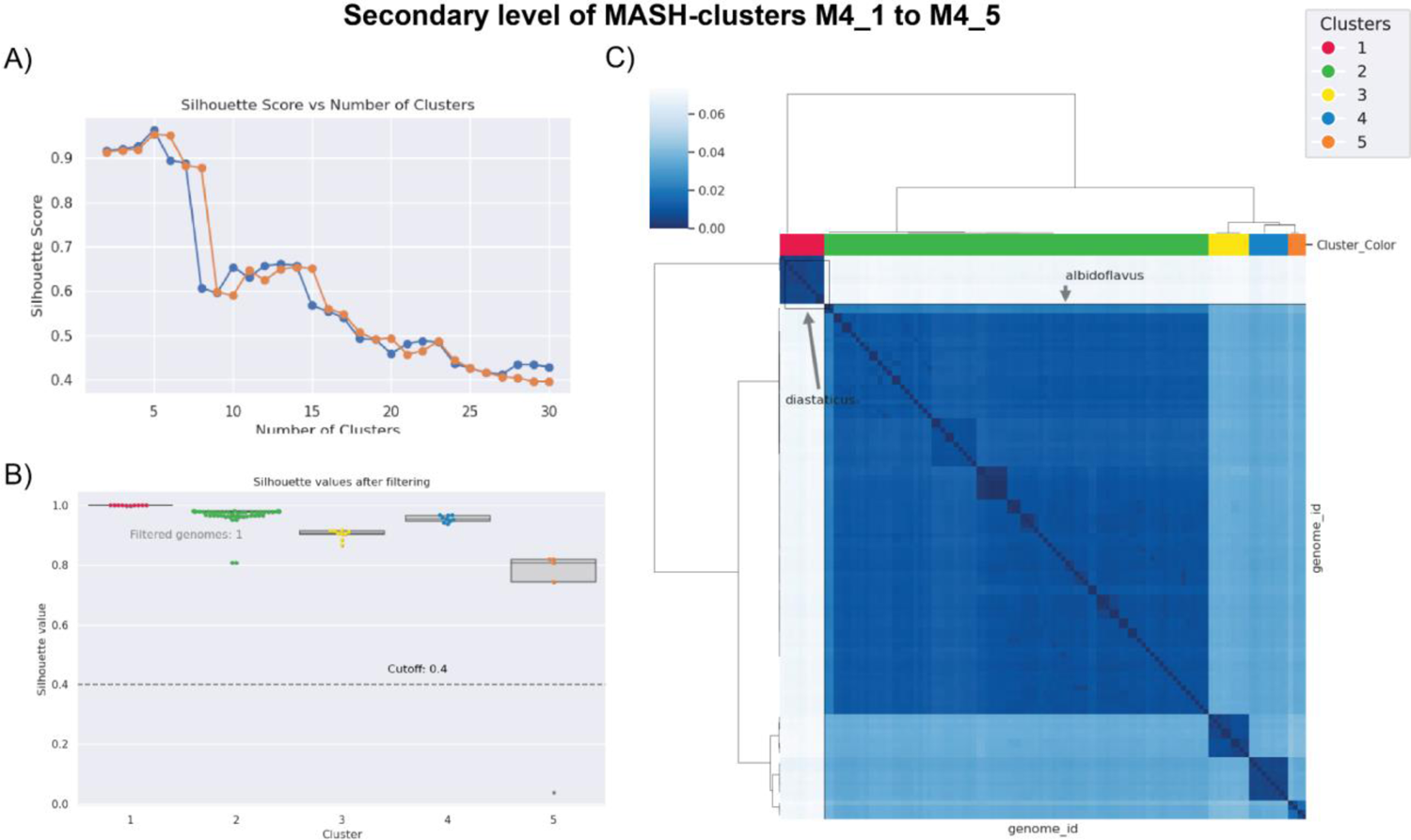
Detection of secondary MASH-clusters using Silhouette scores within the M4 primary MASH-cluster A) The average silhouette scores of all samples against the number of defined clusters with hierarchical clustering based on the MASH distance matrix. The orange line plot represents the original dataset of M4 MASH-cluster genomes whereas the blue represents the dataset after removing poorly clustered samples. B) The silhouette scores of each sample across 5 secondary MASH-clusters. The cutoff of 0.4 was used to select the samples with good clustering. The grey dots represent 1 genome that was removed from the clustering analysis. C) Heatmap representing the MASH distances between the genomes from the refined dataset. The rows and columns are clustered using the hierarchical clustering method where the colors on columns represent the 5 secondary MASH-clusters. The highlighted text on the heatmap represents some of the abundant species including *S. albidoflavus* as a major contributor of the M4 MASH-cluster.

**Figure S10.**
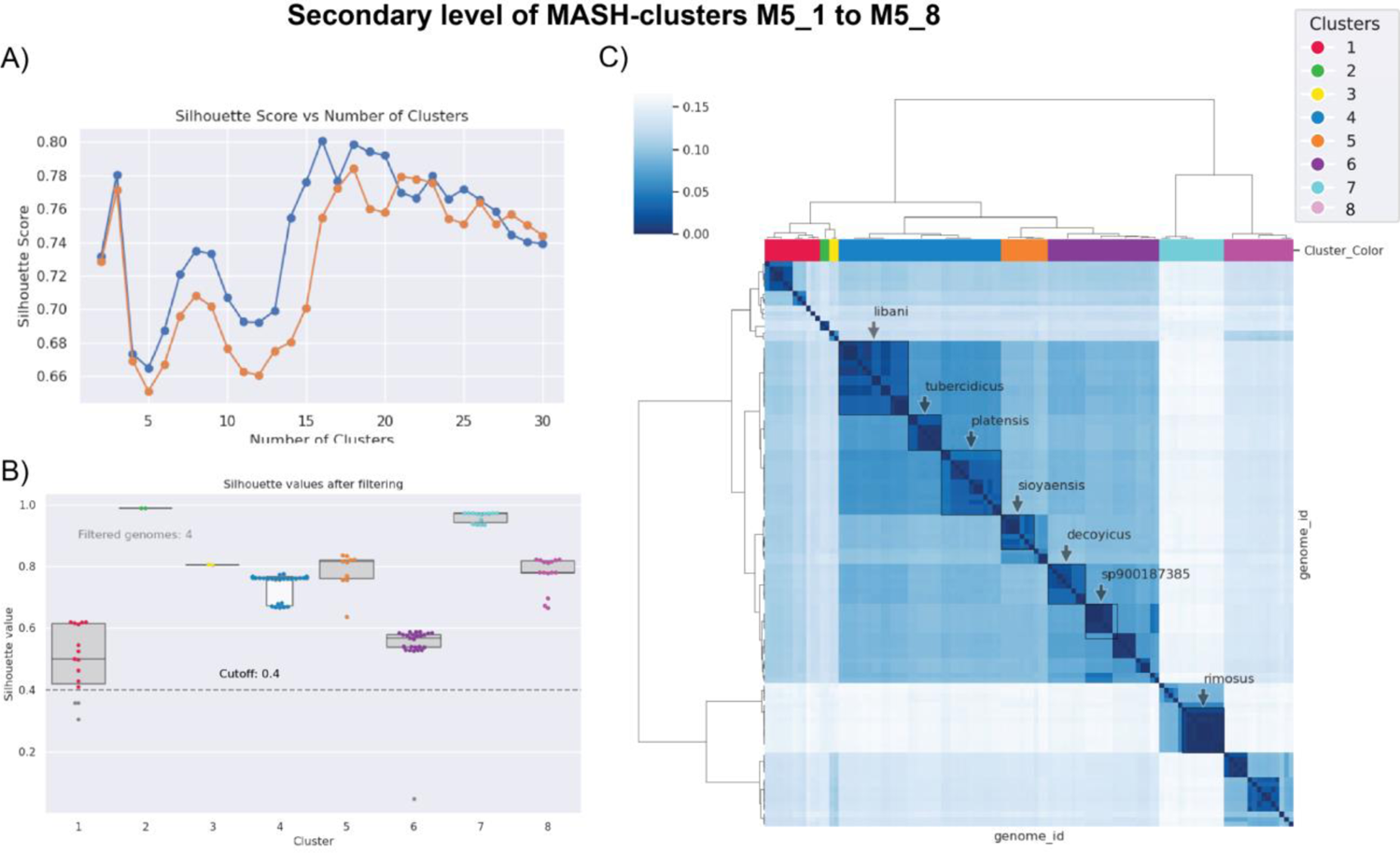
Detection of secondary MASH-clusters using Silhouette scores within the M5 primary MASH-cluster A) The average silhouette scores of all samples against the number of defined clusters with hierarchical clustering based on the MASH distance matrix. The orange line plot represents the original dataset of M5 MASH-cluster genomes whereas the blue represents the dataset after removing poorly clustered samples. B) The silhouette scores of each sample across 8 secondary MASH-clusters. The cutoff of 0.4 was used to select the samples with good clustering. The grey dots represent 4 genomes that were removed from the clustering analysis. C) Heatmap representing the MASH distances between the genomes from the refined dataset. The rows and columns are clustered using the hierarchical clustering method where the colors on columns represent the 8 secondary MASH-clusters. The highlighted text on the heatmap represents some of the abundant species. Note that the M5 MASH-cluster is one of the most diverse and likely poorly sampled in the dataset.

**Figure S11.**
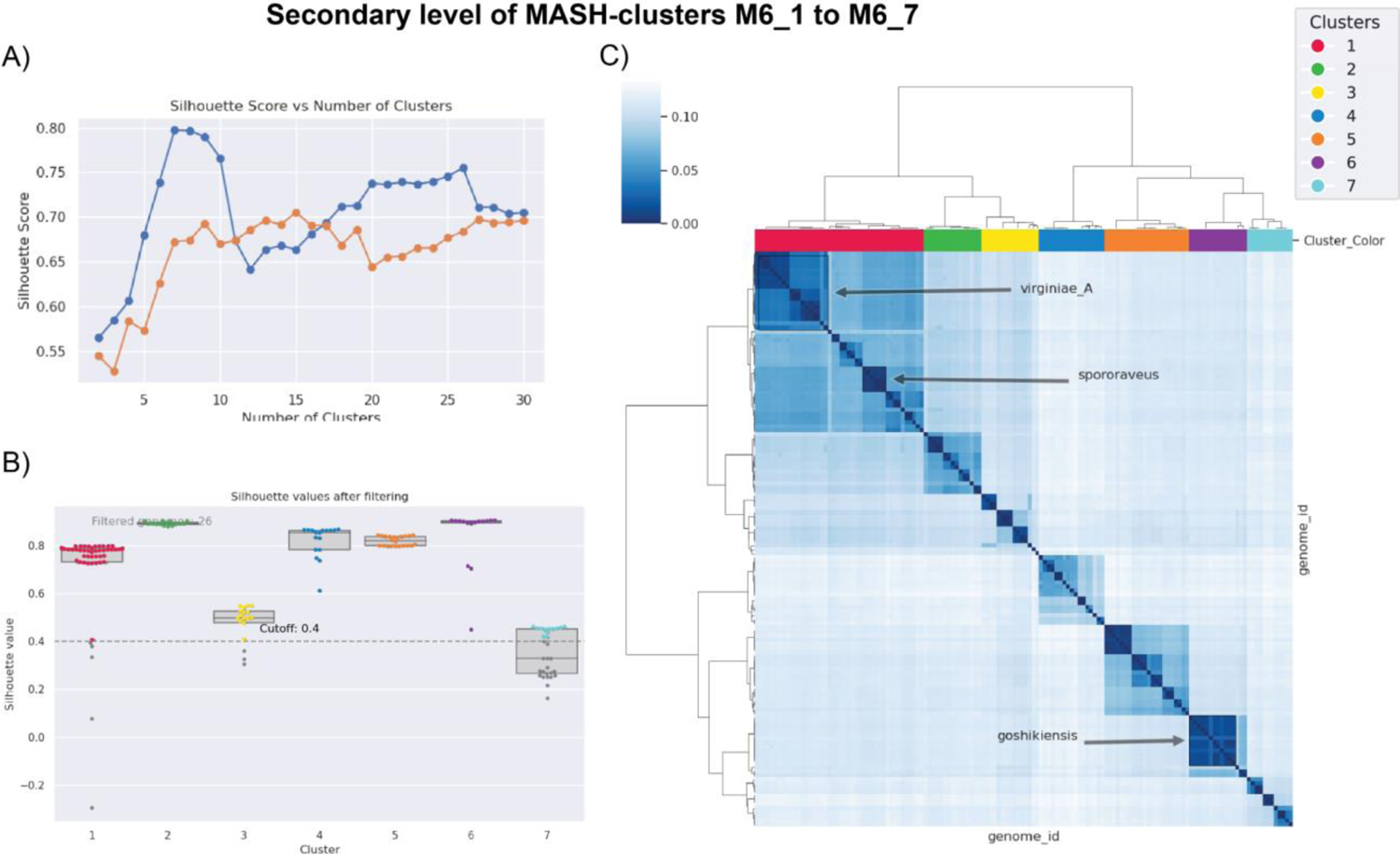
Detection of secondary MASH-clusters using Silhouette scores within the M6 primary MASH-cluster A) The average silhouette scores of all samples against the number of defined clusters with hierarchical clustering based on the MASH distance matrix. The orange line plot represents the original dataset of M6 MASH-cluster genomes whereas the blue represents the dataset after removing poorly clustered samples. B) The silhouette scores of each sample across 7 secondary MASH-clusters. The cutoff of 0.4 was used to select the samples with good clustering. The grey dots represent 26 genomes that were removed from the clustering analysis. C) Heatmap representing the MASH distances between the genomes from the refined dataset. The rows and columns are clustered using the hierarchical clustering method where the colors on columns represent the 7 secondary MASH-clusters. The highlighted text on the heatmap represents some of the abundant species.

**Figure S12.**
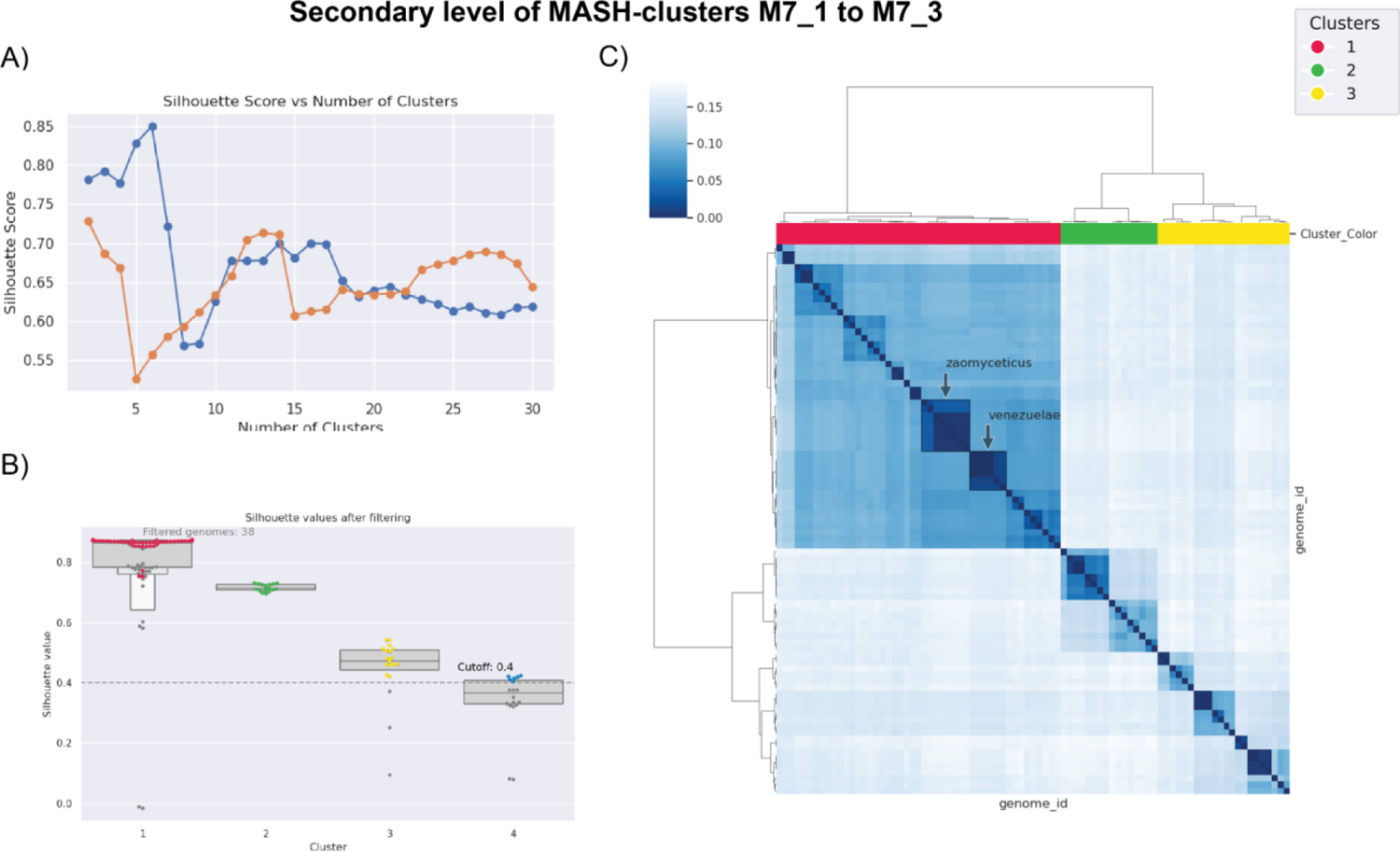
Detection of secondary MASH-clusters using Silhouette scores within the M7 primary MASH-cluster A) The average silhouette scores of all samples against the number of defined clusters with hierarchical clustering based on the MASH distance matrix. The orange line plot represents the original dataset of M7 MASH cluster genomes whereas the blue represents the dataset after removing poorly clustered samples. B) The silhouette scores of each sample across 4 secondary MASH-clusters. The cutoff of 0.4 was used to select the samples with good clustering. The grey dots represent 38 genomes that were removed from the clustering analysis. C) Heatmap representing the MASH distances between the genomes from the refined dataset (note that 3 clusters were generated in the refined dataset). The rows and columns are clustered using the hierarchical clustering method where the colors on columns represent the 3 secondary MASH-clusters. The highlighted text on the heatmap represents some of the abundant species.

**Figure S13.**
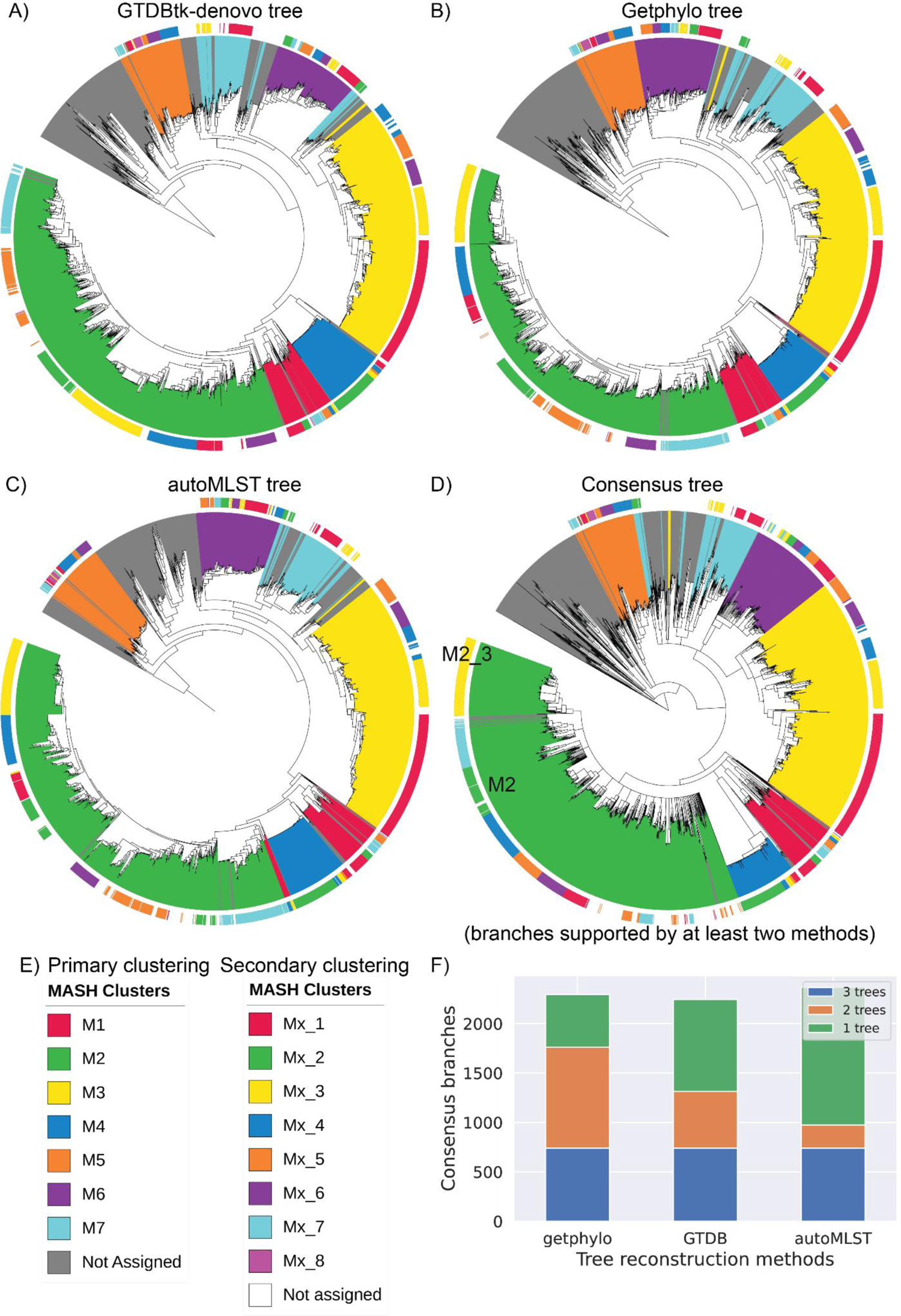
Comparative assessment of MASH clusters aligned against different phylogenetic trees Phylogenetic trees were reconstructed using 3 different methods: A) GTDB-Tk denovo, B) getphylo, and C) autoMLST. D) A consensus tree generated from the getphylo tree where the branches supported in at least two trees were kept. E) Color legend representing the primary and secondary level of MASH clusters. For example, M2 and M2_3 are highlighted in panel D. F) Number of branches in individual trees showing the consensus across the other trees, with getphylo showing maximum number of consensus branches.

**Figure S14:**
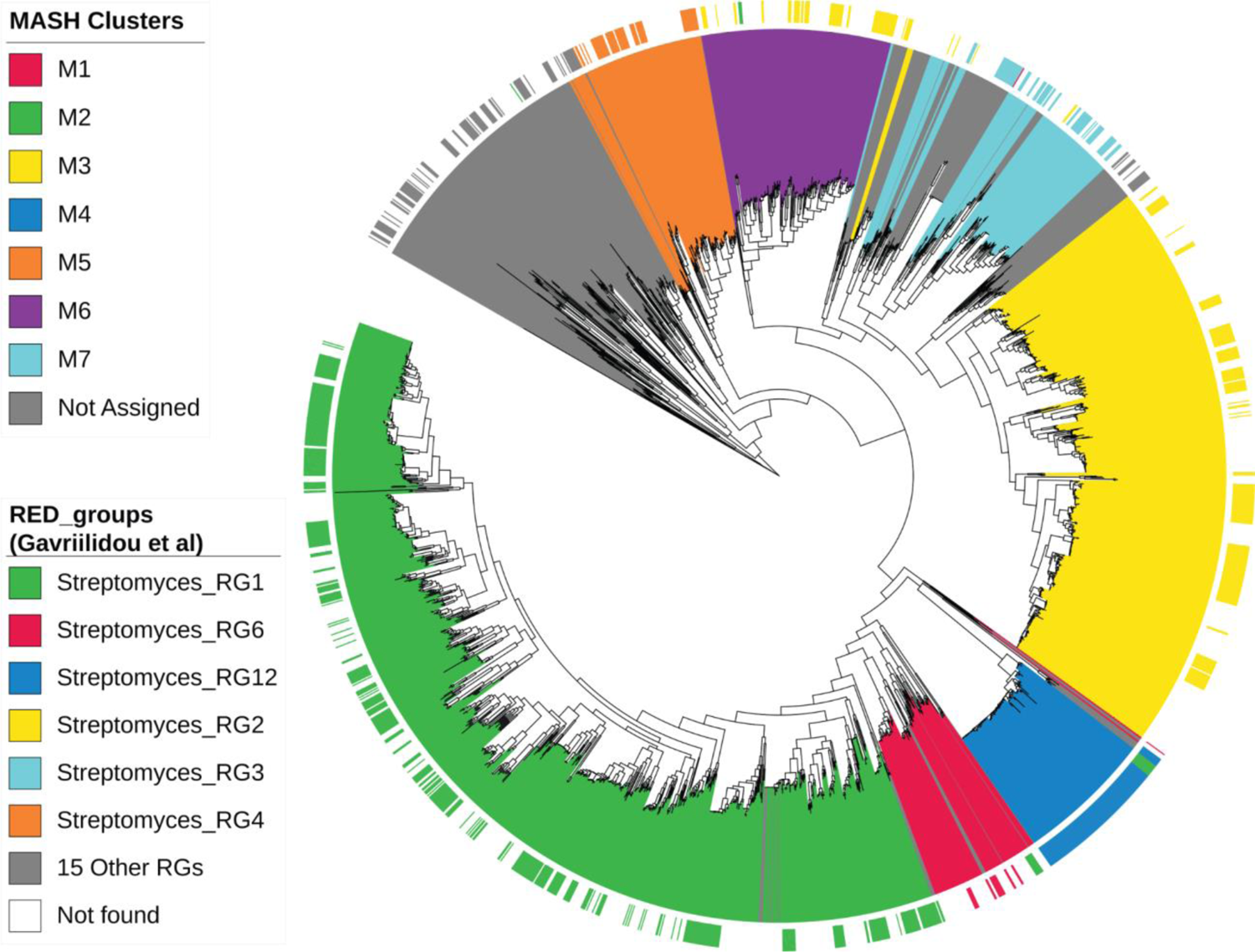
Comparison of MASH-clusters with groups proposed by Gavriilidou et. al. The RED (relative evolutionary divergence) groups as defined by Gavriilidou et. al.[2] were mapped to the consensus tree and the MASH-clusters. The top 6 RED_groups are represented by different colors on the external strip with the remaining RED_groups colored in grey. The GTDB species that were not part of the earlier study are ignored on color color strip.

**Figure S15.**
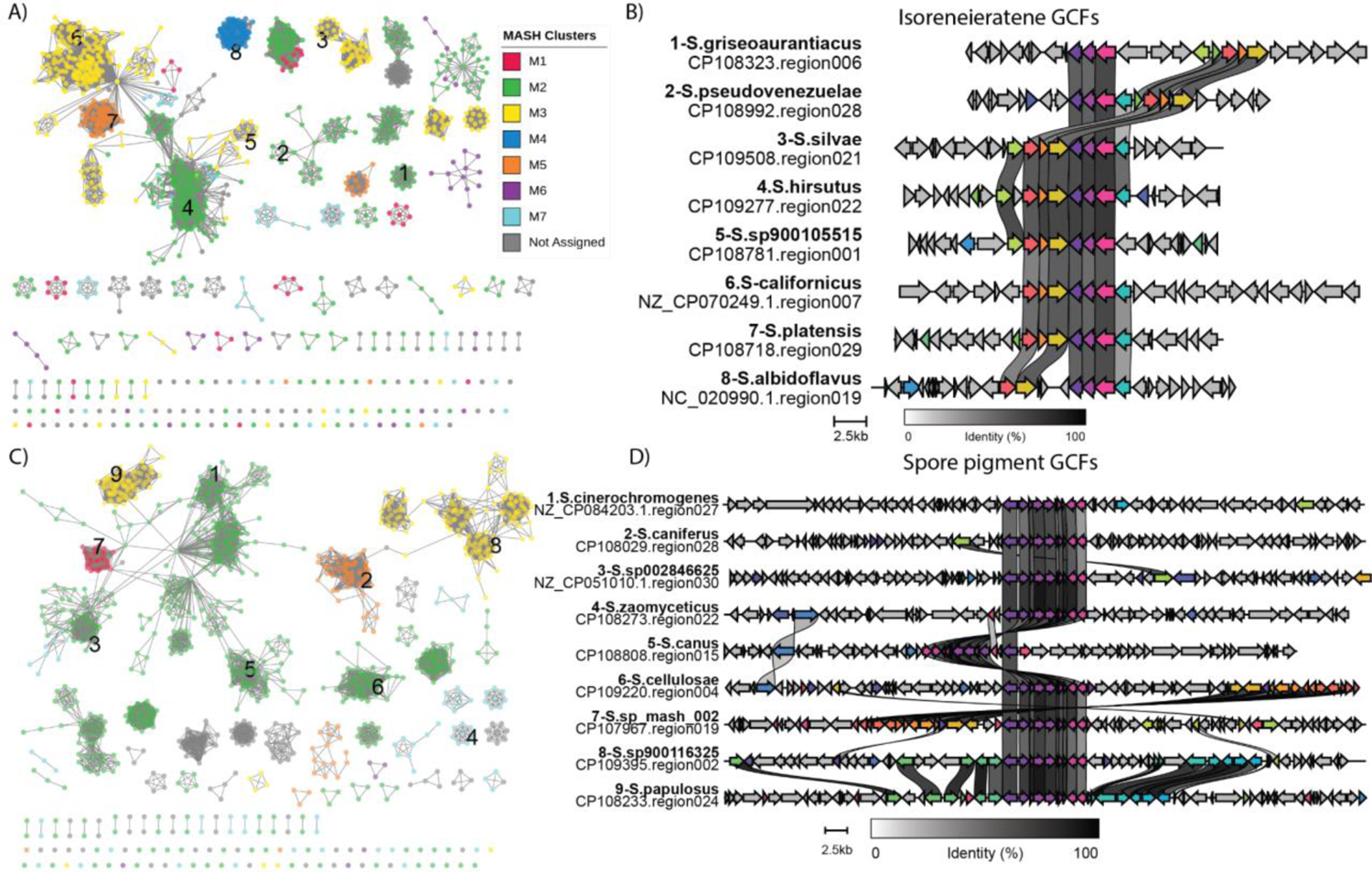
Distribution and variation within common GCFs. A) Similarity network based on BiG-SCAPE indicating detection of different GCFs for BGCs with hits against a common MIBiG entry of isorenieratene. B) The alignment of selected BGCs from panel A (highlighted with numbers) indicates mostly conserved core biosynthetic genes with variations arising from extended cluster boundary definitions. C) Similarity network based on BiG-SCAPE indicating detection of different GCFs for BGCs with hits against a common MIBiG entry of spore pigment. D) The alignment of selected BGCs from panel C (highlighted with numbers) again indicating highly conserved core biosynthetic genes with variations arising from extended cluster boundary definitions.

**Figure S16.**
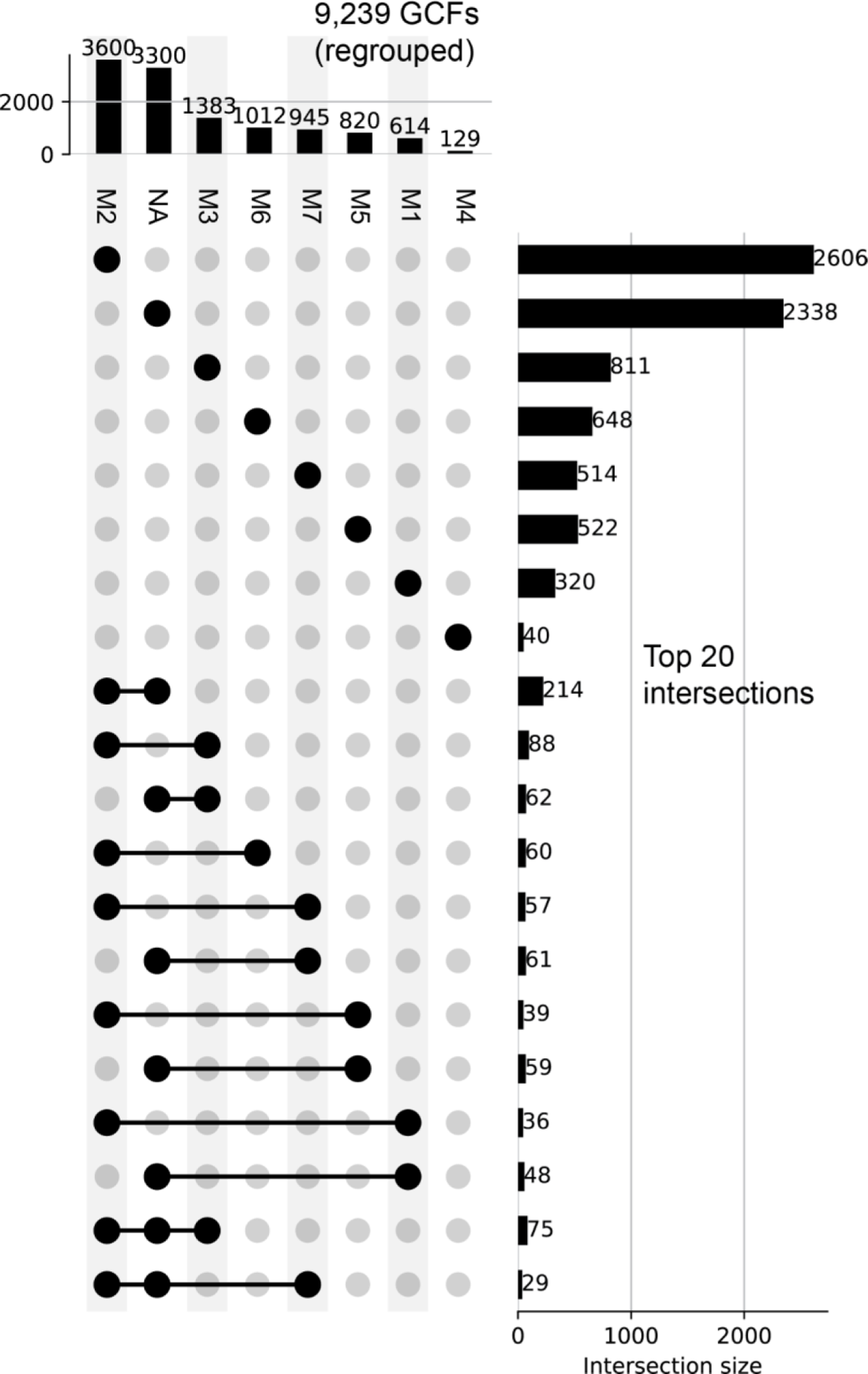
UpSet plot representation of GCFs present across different primary MASH-clusters The ‘*NA’* category is represented by genomes with no MASH-cluster assigned. The top 20 most abundant intersections are selected for the visualization. The bars along the top represent total GCFs present in each MASH-cluster. The bars along the right represent the number of GCFs in the corresponding intersection.

**Figure S17:**
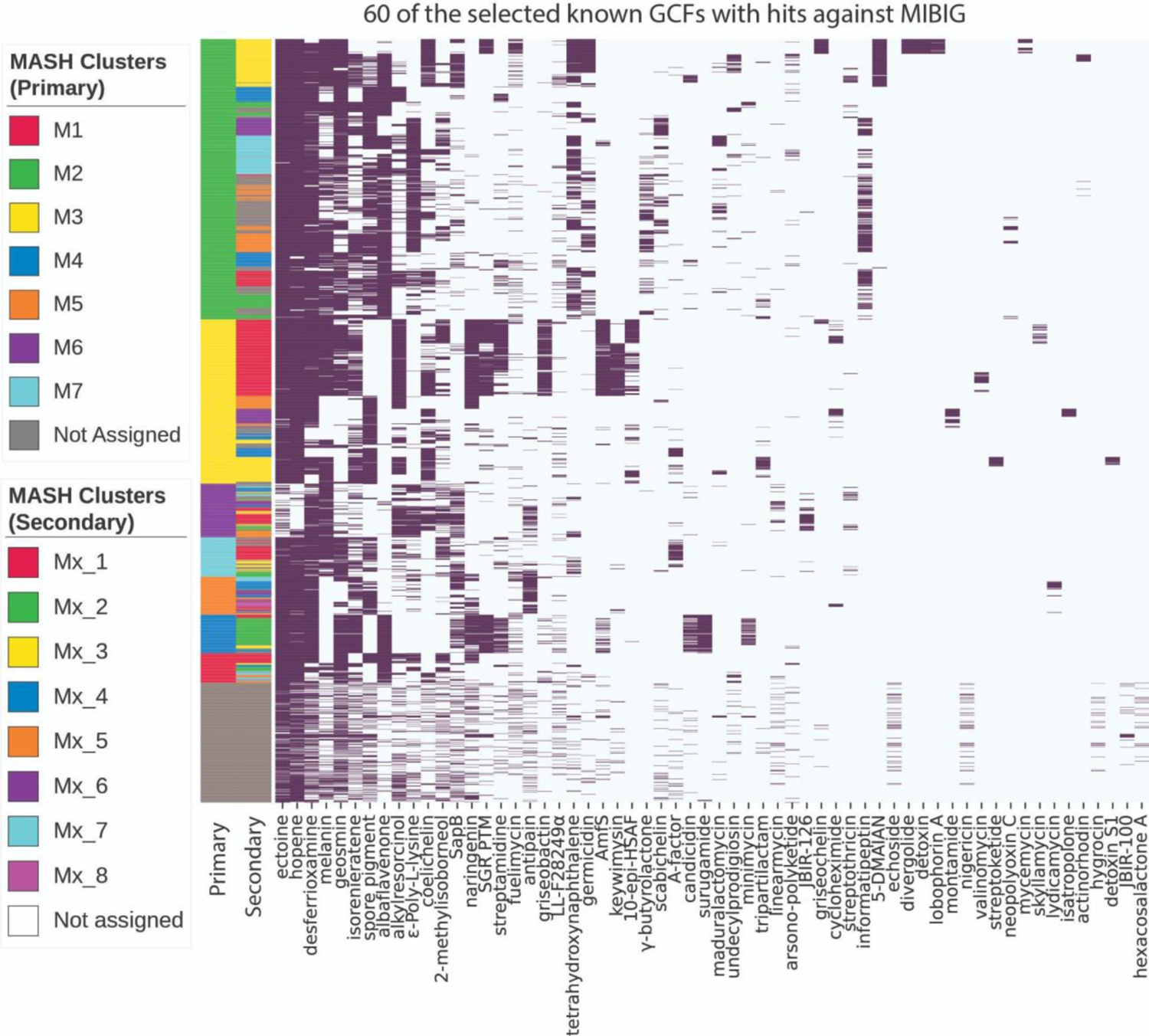
The presence-absence heatmap of GCFs with knownclusterblast similarity hits against the MIBiG database. The top 20 GCFs were selected from each of the three categories: present in more than 6 MASH-clusters, present in 2 to 6 MASH-clusters, and present in only one of the MASH-clusters. The row colors represent MASH-cluster assignment at both primary and secondary levels. The secondary MASH-cluster colors are assigned within each primary MASH-cluster. For example, M4_1 to M4_5 are assigned the colors of Mx_1 to Mx_5 in the legend for secondary MASH-clusters.

**Figure S18.**
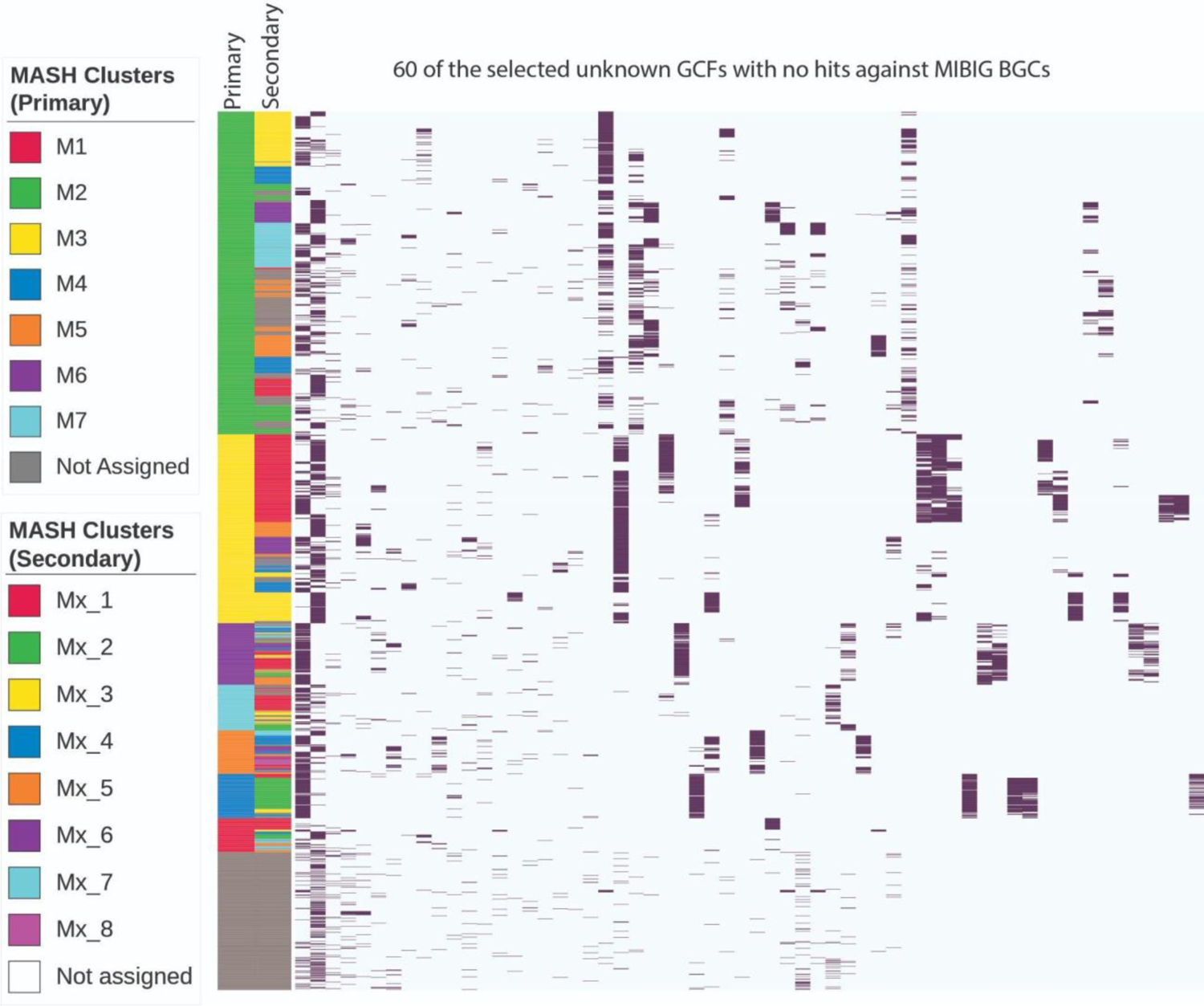
The presence-absence heatmap of GCFs without knownclusterblast similarity hits against the MIBiG database. The top 20 GCFs were selected from each of the three categories: present in more than 6 MASH-clusters, present in 2 to 6 MASH-clusters, and present in one of the MASH-clusters. The row colors represent MASH-cluster assignment at both primary and secondary levels. The secondary MASH-cluster colors are assigned within each primary MASH-cluster. For example, M4_1 to M4_5 are assigned the colors of Mx_1 to Mx_5 in the legend for secondary MASH-clusters.

**Figure S19.**
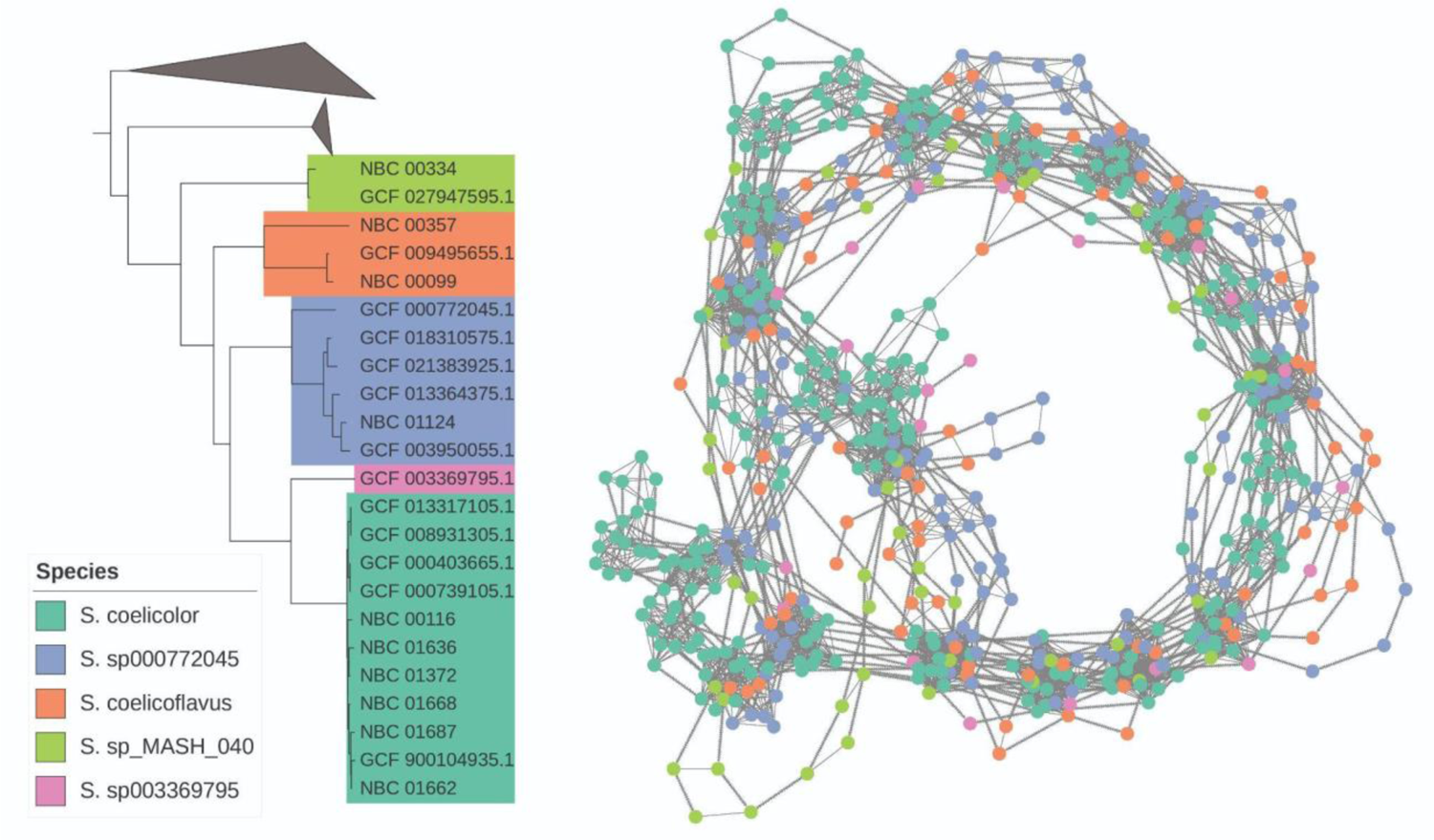
Similarity network integrated with chromosomal order of BGCs across multiple species (*Left*) Phylogenetic tree representing 23 genomes belonging to M2_3 secondary MASH-cluster. The other clades of the MASH-cluster were collapsed. (*Right*) Similarity network integrating chromosomal order across 5 different species of MASH-cluster M2_3 depicting the conserved and variable BGCs across the genomes.

